# A stable microtubule bundle formed through an orchestrated multistep process controls quiescence exit

**DOI:** 10.1101/2023.05.02.539064

**Authors:** Damien Laporte, Aurélie Massoni-Laporte, Charles Lefranc, Jim Dompierre, David Mauboules, Emmanuel. T. Nsamba, Anne Royou, Lihi Gal, Maya Schuldiner, Mohan L. Gupta, Isabelle Sagot

**Affiliations:** Univ. Bordeaux, CNRS, IBGC, UMR 5095, Bordeaux, France; Genetics, Development, and Cell Biology, Iowa State University, Ames, IA 50011, USA.; Department of Molecular Genetics, Weizmann Institute of Science, Rehovot, Israel

## Abstract

Cells fine-tune microtubule assembly in both space and time, to give rise to distinct edifices with specific cellular functions. In proliferating cells, microtubules are highly dynamics, and proliferation cessation often leads to their stabilization. One of the most stable microtubule structures identified to date is the nuclear bundle assembled in quiescent yeast. In this report, we characterize the original multistep process driving the assembly of this structure. This Aurora B-dependent mechanism follows a precise temporality that relies on the sequential actions of kinesin-14, kinesins-5 and involves both microtubule-kinetochore and kinetochore-kinetochore interactions. Upon quiescence exit, the microtubule bundle is disassembled via a cooperative process involving kinesin-8 and its full disassembly is required prior to cells re-entry into proliferation. Overall, our study provides the first description, at the molecular scale, of the entire life cycle of a stable microtubule structure *in vivo*, and sheds light on its physiological function.

## Introduction

Microtubules (MTs) are hollow cylindrical polymers assembled by the non-covalent interaction of α- and β-tubulin heterodimers. MTs are generally nucleated at MT organizing centers (MTOC) by a ɣ-tubulin complex (ɣ-TuC) which acts as a template and stabilizes the so-called MT “minus-end” (Liu et al., 2021; Roostalu and Surrey, 2017; Sanchez and Feldman, 2017; Thawani and Petry, 2021). At the opposite end, the “plus-end” (+end), MTs elongate by the addition of GTP tubulin. The β-tubulin bound GTP is hydrolyzed and stable but transient GDP + Pi intermediates are generated in the region immediately behind the growing +end. Subsequent Pi release favors MT depolymerization which can be rescued by *de novo* GTP-tubulin addition (Cleary and Hancock, 2021; Gudimchuk and McIntosh, 2021). Thus, MTs alternate between periods of growth and shrinkage, a behavior termed dynamic instability (Mitchison and Kirschner, 1984). *In vivo*, a profusion of MT-associated proteins (MAPs) regulate MT length and dynamics (Bodakuntla et al., 2019; Goodson and Jonasson, 2018). In addition, MTs are often assembled from multiple tubulin variants or isotypes, and are modified by a cohort of post-translational modifications that modulate MT dynamics directly or by influencing the recruitment of MAPs (Janke and Magiera, 2020; Roll-Mecak, 2020). Other MAPs organize MTs into high-ordered assemblies, either by cross-linking MTs or by connecting them with cellular structures such as membranes, chromosomes or actin cytoskeleton components (Bodakuntla et al., 2019; Meiring et al., 2020). Cells fine-tune these mechanisms in both space and time to give rise to distinct MT edifices with specific functions.

In proliferating cells, MT dynamics are crucial for their functions. It allows exploration of the cell volume in search of structures to “capture”, such as centromeres during mitosis, or to exert pushing or pulling forces required for various cellular processes such as cell migration or morphogenesis (Heald and Khodjakov, 2015; Kirschner and Mitchison, 1986). The MT cytoskeleton can undergo extensive rearrangements and assemble more stable MT structures, especially when cells change fate (Meiring et al., 2020; Röper, 2020). For example, in terminally differentiated cells, such as epithelial cells, cardiomyocytes or neurons, stable MT networks make up the majority, and ensure critical cellular functions such as cell shape maintenance or long-distance intracellular transport (Baas et al., 2016; Muroyama and Lechler, 2017). Defects in MT stabilization are at the origin of many human pathologies, including neurodegenerative diseases and ciliopathies (Anvarian et al., 2019; Wheway et al., 2018). Effectors responsible for MT stabilization have been actively searched for since the 1980’s. In mammals, the involvement of specific MAPs such as Tau, MAP2, MAP6 and PRC1, or tubulin post-translational modifications has been extensively studied. While their contributions to MT stabilization are central and undisputed, their sole actions do not fully explain the observed levels of MT stability in many types of non-dividing cells (Hahn et al., 2019).

For years, yeast species have been instrumental in unraveling mechanisms that regulate MT dynamics in eukaryotes. While proliferating yeast cells exhibit a dynamic MT network (Winey and Bloom, 2012), proliferation cessation goes with the formation of dramatically stable MT structures in both *S. cerevisiae* and *S. pombe* (Laporte and Sagot, 2014; Laporte et al., 2013, 2015). Upon quiescence establishment, *S. cerevisiae* cells assemble a bundle composed of stable parallel MTs, hereafter referred to as the Q-nMT bundle, for <u>Q</u>uiescent-cell <u>n</u>uclear <u>M</u>icro<u>t</u>ubule bundle. This structure originates from the nuclear side of the spindle pole body (SPB), the yeast equivalent of the centrosome. It spans the entire nucleus and relocalizes kinetochores and centromeres, which remain attached to MT +ends (Laporte and Sagot, 2014). Because the MTs embedded in the Q-nMT bundle are not all of the same length, the structure has a typical arrow shape. When cells exit quiescence, the Q-nMT bundle depolymerizes and, by pulling the attached centromeres back to the SPB, allows the recovery of the typical Rabl-like configuration of chromosomes found in G1 yeast cells (Jin et al., 1998). The molecular mechanisms underlying the formation of this peculiar stable MT structure and its physiological function(s) are not understood. However, cells defective in Q-nMT bundle formation have a compromised survival in quiescence and a drastically reduced fitness upon cell cycle re-entry (Laporte and Sagot, 2014; Laporte et al., 2013, 2015).

Here, we show that Q-nMT bundle formation is a multistep process that follows a precise temporal sequence. The first step relies on Aurora B/Ipl1 activity and requires the kinesin-14 Kar3, its regulator Cik1, and the EB1 homolog Bim1. It leads to the formation of a short (≈ 1 µm) and stable bundle that resembles a half mitotic spindle. Importantly, in this first step, MT polymerization and stabilization are coupled. In a second step, additional MTs polymerize from the SPB, elongate, and are zipped to pre-existing MTs in a Cin8/kinesin-5 dependent manner. Complete stabilization of the Q-nMT bundle is achieved in a third and final step that requires the kinesin-5 Kip1. Our observations further indicate that Q-nMT bundle assembly requires both MT-kinetochore attachment and inter-kinetochore interactions. Finally, we show that, upon exit from quiescence, the Q-nMT bundle disassembles from its +ends, each MT depolymerizing in coordination with its neighbors, via the action of the depolymerase Kip3, a member of the kinesin-8 family. Importantly, we show that the complete disassembly of the Q-nMT bundle is required for cells to re-enter the proliferation cycle. Overall, this study describes the entire life cycle of a MT structure, from the molecular mechanisms involved in its formation and stabilization to its disassembly and further suggests that this atypical quiescent specific structure could act as a regulator, or a “control point” for cell cycle resumption upon exit from quiescence.

## Results

### The Q-nMT bundle formation is a three-step process

We have previously shown that the Q-nMT bundle observed in quiescent cells is a stable nuclear structure (Laporte et al., 2013). To investigate whether the stabilization of the Q-nMT bundle was concomitant with its assembly or whether it assembled first as a dynamic structure and then gets stabilized, we quantified the Q-nMT bundle length (Fig. 1A) and thickness (Fig. 1B) upon quiescence establishment following carbon source exhaustion. We found that the Q-nMT bundle assembly process could be divided into 3 sequential phases. In an initial phase (phase I), MTs elongated from the SPB to reach ≈ 0.8 µm. The number of these MTs, referred to as phase I-MTs, was approximately the same as in a mitotic spindle (see inset of Fig. 1B and Sup. Fig. 1A). Importantly, phase I-MTs were resistant to nocodazole (Noc), a MT poison that causes dynamic MT depolymerization, indicating that phase I-MT stabilization was concomitant with their polymerization (Fig. 1A-B). In a second phase, beginning ≈ 10h after glucose exhaustion, additional MTs emerged from the SPB (phase II-MTs) and elongated along phase I-MTs, nearly doubling the thickness of the phase I MT bundle (Fig. 1B and Sup Fig. 1A). At this stage, the tip of the newly elongated MTs were unstable (Fig. 1A-B). Complete stabilization of the Q-nMT bundle was achieved ≈ 48h after glucose exhaustion (phase III). Indeed, after this step, a Noc treatment did not affect either the length or the thickness of the Q-nMT bundle (Fig. 1A-B and Sup. Fig. 1B). Plotting Q-nMT bundle width as a function of length for each individual cell before and after Noc treatment further revealed the 3 successive phases of the Q-nMT bundle formation (Fig. 1C). Of note, Q-nMT bundles were also observed both in diploid cells upon glucose exhaustion (Sup. Fig. 1C) and in haploid cells transferred to water (Sup. Fig. 1D).

**Figure 1:**
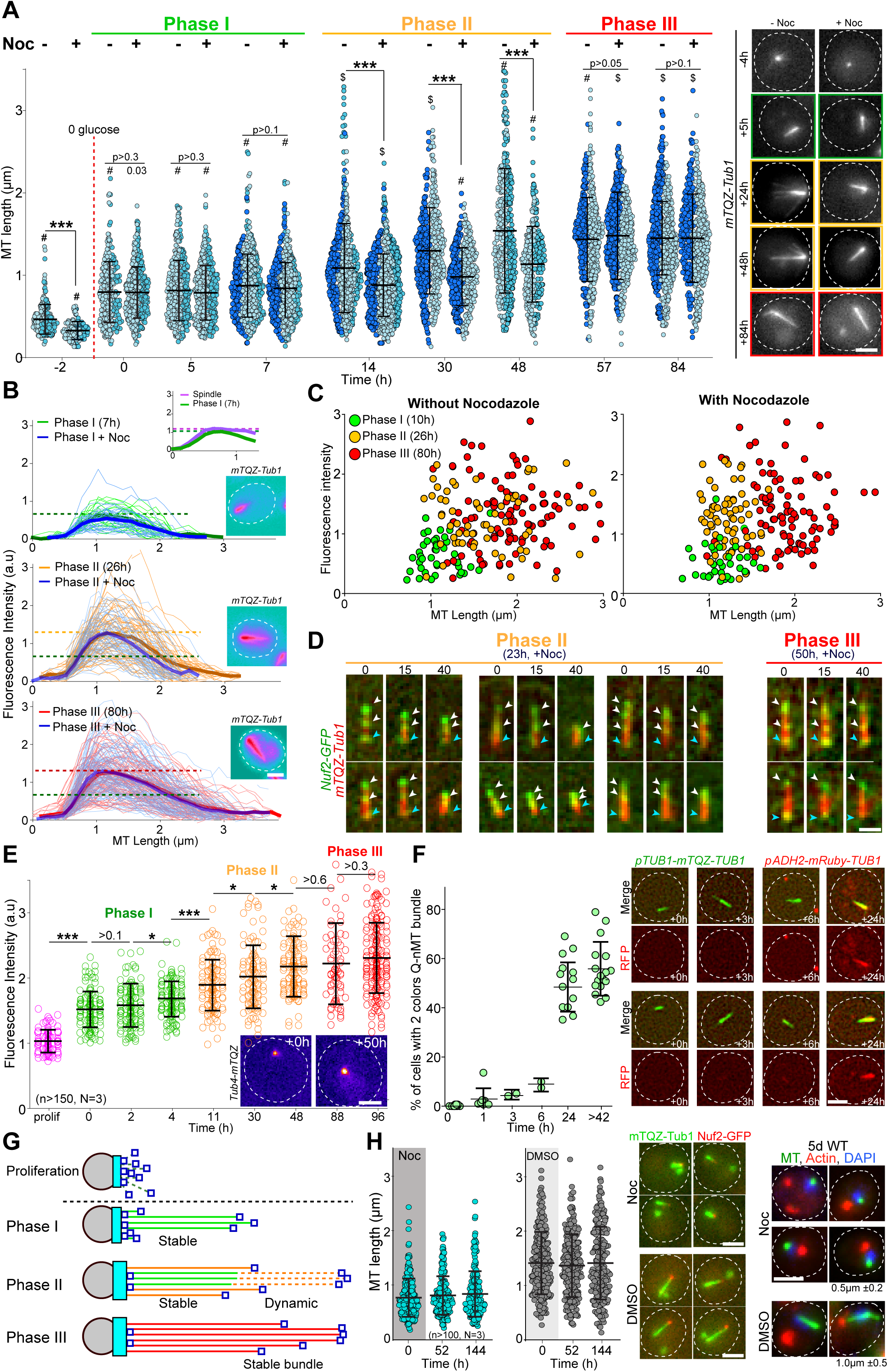
The formation of the Q-nMT bundle is a three-step process. **(A)** Nuclear MT length in WT cells expressing mTQZ-Tub1, before (**-**) or after (**+**) a 15 min Noc treatment (30 µg/ml) upon entry into quiescence. Each circle corresponds to the length of an individual MT structure. Three independent experiments are shown (pale blue, cyan and dark blue, n > 160 for each point in each experiment). The mean and SD are shown. Student test or ANOVA (sample >2) were used to compare inter-replicates. #: p-value > 0.05, $: p-value < 0.05. A student test (t-test with two independent samples) was used to compare the results obtained with or without Noc, the indicated p-values being the highest measured among experiments. ***: p-value < 1.10^-5^. Images of representative cells are shown. Bar is 2 µm. **(B)** MT fluorescence intensity as a proxy of MT structure width in WT cells expressing mTQZ-Tub1. Mean intensity measurement for half pre-anaphase mitotic spindles (purple), phase I (green), phase II (orange) or phase III (red) Q-nMT bundle. A line scan along the MT structure for individual cells are shown as thin lines, the mean as a bold line (n > 60 / phase), all the lines being aligned at 0,5 µm before the fluorescence intensity increase onset on the SPB side. The blue lines are results obtained after a 15 min Noc treatment (30 µg/ml). In each graph, horizontal dashed line indicates the mean intensity. Images in pseudo-colors of a representative cell for each phase are shown. Bar is 2 µm. **(C)** MT bundle length as a function of MT bundle width for individual cells in each phase before and after a 15 min Noc treatment (30 µg/ml) in WT cells expressing mTQZ-Tub1. Each circle represents an individual MT structure. **(D)** WT cells expressing mTQZ-Tub1 (red) and Nuf2-GFP (green) in phase II (23 h) or phase III (50 h) were deposited on an agarose pad containing 30 µg/ml Noc and imaged. Blue arrowheads: SPB; white arrowheads: Nuf2-GFP clusters. Time is in min after deposition on the pad. Bar is 1 µm. **(E)** Tub4-mTQZ fluorescence intensity measured at the SPB upon entry into quiescence. Each circle represents an individual cell. The mean and SD are shown; t-tests were used to compare independent samples (N = 3, n > 150), *: 0.05 > p-value > 1.10^-3^, ***: p-value < 1.10^-5^. Images in pseudo-colors of representative cells are shown. Bar is 2 µm. **(F)** WT cells expressing mTQZ-Tub1 under the *TUB1* promoter and mRuby-Tub1 under the *ADH2* promoter. The percentage of cells harboring both mTQZ and mRuby fluorescence along the Q-nMT bundle is shown; each circle being the percentage for an independent experiment, with n > 200 cells counted for each experiment. The mean and SD are shown. Images of representative cells at the indicated time after glucose exhaustion are shown. Bar is 2 µm. **(G)** Schematic of the Q-nMT bundle formation. During phase I, stable MTs (phase I-MT - green) elongate from the SPB (grey). During phase II, the amount of Tub4 (cyan) increases at the SPB. In the meantime, new MTs (phase II-MT - orange) elongated from the SPB and are stabilized along the phase I-MTs, yet their +ends remain dynamic (dashed lines). After phase III, all MTs are stabilized (red). Nuf2 is schematized as dark blue squares. **(H)** Upon glucose exhaustion, WT cells expressing mTQZ-Tub1 (green) and Nuf2-GFP (red) were pulsed treated with 30 µg/ml Noc (blue) or DMSO (grey) for 24 h. Noc or DMSO were then chased using carbon exhausted medium and cells were imaged. Each circle corresponds to MT structure length in an individual cell. The mean and SD are shown (N = 3, n > 100). Images of representative cells 2 d after the chase and representative cells 5 days after the chase are shown. For the right panels, tubulin (green) was detected by immunofluorescence, actin (red) by phalloidin and DNA (blue) with DAPI. The mean Q-nMT bundle length (±SD) in the population is indicated.

To confirm the above observations, first, we followed Nuf2-GFP, a protein that localizes to the MT +end. As expected, movies showed that upon Q-nMT bundle formation initiation, the Nuf2-GFP signal relocated from the SPB to the distal part of elongating Q-nMT bundles (Sup. Fig. 1E). In phase II, when cells were treated with Noc, the Nuf2-GFP signal followed depolymerizing MTs (Fig. 1D, left panel), testifying for the instability of phase II-MTs. In contrast, the Nuf2-GFP signal remained immobile upon Noc treatment in phase III, as cells have fully stabilized Q-nMT bundles (Fig. 1D, right panel). Second, we measured the γ-tubulin (Tub4) signal at the SPB and found that Tub4 started to accumulate at the SPB at the beginning of phase I to reach a plateau at the end of phase II (Fig. 1E and Sup. Fig. 1F). Interestingly, the amount of Tub4 at the SPB was proportional to the thickness of the Q-nMT bundle (Sup. Fig. 1G). Finally, in order to confirm the two waves of MT elongation, we developed a strain expressing mTQZ-Tub1 from the endogenous TUB1 promoter and mRuby-TUB1 under the ADH2 promoter, a promoter that is de-repressed after glucose exhaustion, as confirmed by western blot (Sup. Fig. 1H). We observed that mRuby-TUB1 was incorporated into the Q-nMT bundle only after phase I (Fig. 1F and Sup. Fig. 1H) confirming the existence of a second wave of MT elongation during phase II. All the above experiments led us to propose the model shown in figure 1G.

### Q-nMT bundle formation is an essential time-regulated process

Increasing cytoplasmic viscosity has been shown to dampen MT dynamics (Molines et al., 2022). Upon quiescence establishment, yeast cells undergo a transition from a fluid to a solid-like phase (Munder et al., 2016; Joyner et al., 2016) and an acidification (Jacquel et al., 2021) which could contribute to the formation of the Q-nMT bundle. Several lines of evidence suggest that these changes in physicochemical properties are not involved in Q-nMT formation. First, 4-day-old quiescent cells can simultaneously display both dynamic cytoplasmic MTs (cMTs) and a stable Q-nMT bundle (Sup. Fig. 1I and (Laporte et al., 2013)). Second, when we mimicked a fluid to solid-like phase transition in proliferating cells by artificially lowering the pH, no stable MTs were observed, whereas F-actin aggregation was induced (Sup. Fig. 1J) as expected (Peters et al., 2013). Third, 2-day-old quiescent cells exhibiting a phase III Q-nMT bundle were able to grow *de novo* dynamic cMTs after a pulse-chase Noc treatment (Sup. Fig. 1K).

Importantly, if Q-nMT bundle formation was solely dependent on physicochemical changes, this structure should be able to assemble in late quiescence. By treating cells with Noc upon glucose exhaustion, we were able to prevent the formation of the Q-nMT bundle in early quiescence (Fig. 1H). Strikingly, when the drug was washed out, no Q-nMT bundle assembled, even 144h after Noc removal (Fig. 1H). These cells had entered quiescence because they had assembled actin bodies (Fig. 1H, right panel), another quiescent cell specific structure (Sagot et al., 2006). This experiment strongly suggests that Q-nMT formation is a process induced by a transient signal emitted upon glucose exhaustion.

We have previously shown that mutants unable to assemble Q-nMT bundles have reduced viability in quiescence and a reduced ability to form colonies upon exit from quiescence (Laporte et al., 2013). To establish a direct link between the absence of the Q-nMT bundle and the above phenotypes, we took advantage of our ability to conditionally prevent Q-nMT bundle formation in WT cells using Noc treatment at the onset of glucose exhaustion. As shown in Sup. Fig. 1L, in the absence of the Q-nMT bundle, WT cells lose viability in quiescence and survivors have a reduced ability to generate a progeny upon exit from quiescence. In contrast, a similar Noc treatment of cells that have already assembled a stable Q-nMT bundle (5-day-old cells) did not affect either the viability in quiescence or quiescence exit efficiency, demonstrating that Noc did not have a toxic effect *per se*. Taken together, these experiments confirm our initial findings that the Q-nMT bundle is important for both cell survival during chronological aging and quiescence exit fitness.

### Steady state tubulin levels and Tub3 isotype are critical for Q-nMT bundle formation

We then focused on deciphering the molecular mechanisms involved in Q-nMT bundle formation. Yeast cells express two α-tubulin isotypes, Tub1 and Tub3. In proliferating cells, Tub1 constitutes the majority (Aiken et al., 2019; Nsamba et al., 2021; Schatz et al., 1986). In 4-day-old cells, the two α-tubulin isotypes were found embedded in the Q-nMT bundle. This was observed using two different pairs of fluorescent proteins in order to avoid any influence of the FP variants, and using both widefield and expansion microscopy (Fig. 2A and Sup. Fig. 2A). Since Tub3 stabilizes MTs *in vitro* (Bode et al., 2003), it may be key for Q-nMT bundle formation. To test this hypothesis, we first analyzed the phenotype of *tub3Δ* cells. In phase II, *tub3Δ* cells displayed short MT bundles that were sensitive to Noc. In phase III, no MT bundle were detected (Fig. 2B and Sup. Fig. 2C). Thus, in this mutant, the MT bundles assembled during phase I were not stable and eventually collapsed. In *tub3Δ* cells, the steady state amount of α-tubulin is significantly reduced (Sup. Fig. 2B and (Nsamba et al., 2021)). Thus, the amount of α-tubulin, but not Tub3 itself, may be important for Q-nMT bundle formation. Indeed, mutants defective in α- and β-tubulin folding, such as the prefolding complex mutants *pac10Δ*, *gim3Δ* or *yke2Δ,* or in tubulin heterodimer formation such as *pac2Δ*, or in β-tubulin folding, such as *cin2Δ* or *cin4Δ*, all of which have a reduced amount of tubulin, were unable to assemble a Q-nMT bundle (Sup. Fig. 2D-E).

**Figure 2:**
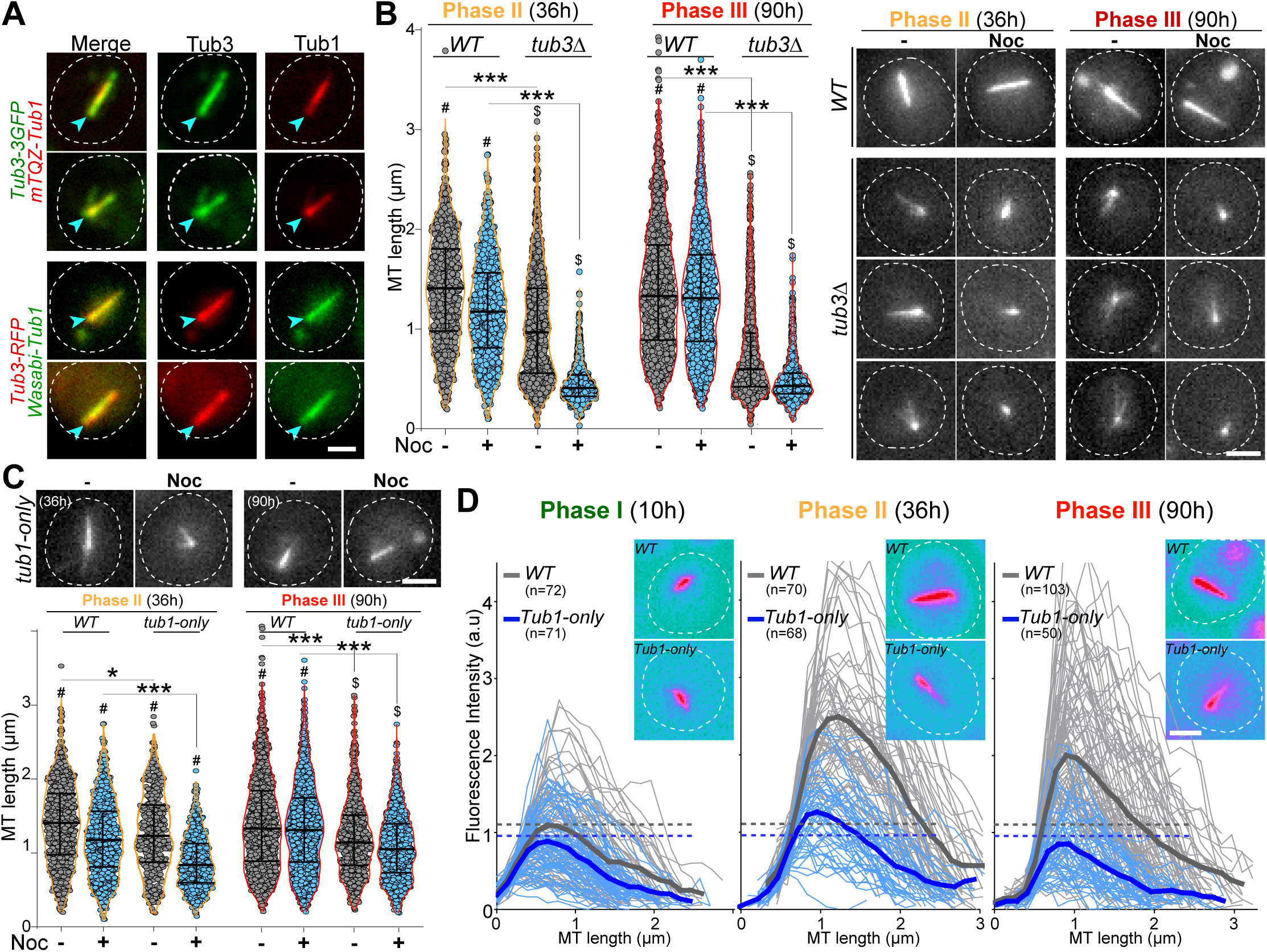
Q-nMT bundle formation is influenced by the alpha-tubulin amount and isotype. **(A)** WT cells (4 d) expressing either Tub3-3GFP (green) and mTQZ-Tub1 (red, top panel) or Tub3-RFP (red) and mWasabi-Tub1 (green, bottom panel). Blue arrowheads point to SPB, bar is 2 µm. **(B)** Nuclear MT length in WT and *tub3Δ* cells expressing mTQZ-Tub1, 36 h (phase II, orange) and 90 h (phase III, red) after glucose exhaustion, treated 15 min (blue) or not (grey) with 30 µg/ml Noc. Each circle corresponds to the length of an individual MT structure, mean and SD are shown. ANOVA was used to compare inter-replicates (n > 200, N = 5); #: p-value > 0.05, $: p-value < 0.05. A student test (t-test with two independent samples) was used to compare (**+**) or (**-**) Noc data. The indicated p-values are the highest calculated among the 5 experiments; ***: p-value < 1.10^-5^. Images of representative cells are shown, bar is 2 µm. **(C)** Nuclear MT length in WT and *Tub1-only* cells expressing mTQZ-Tub1, 36 h (phase II, orange) and 90 h (phase III, red) after glucose exhaustion, treated 15 min or not with 30 µg/ml Noc. Statistical representations are as in (B). Images of representative cells are shown, bar is 2 µm. **(D)** Fluorescence intensity along the Q-nMT bundle in WT (grey) and *Tub1-only* (blue) cells expressing mTQZ-Tub1 grown for 10 h (phase I), 36 h (phase II), and 90 h (phase III). Individual fluorescent intensity is shown as thin line, the mean as a bold line (n > 60 / phase), all the lines being aligned at 0,5 µm before the fluorescence intensity increase onset on the SPB side. Dashed lines indicate the maximal mean fluorescence intensity. Images in pseudo-color of representative cells are shown, bar is 2 µm.

To identify the role of Tub3 *per se*, we took advantage of the “*Tub1-only*” mutant, in which the *TUB3* gene was replaced by the *TUB1* gene at the *TUB3* locus. This mutant expresses only Tub1 and has an α-tubulin level similar to WT (Sup. Fig. 2B and (Nsamba et al., 2021)). We found that “*Tub1-only*” cells have unstable Q-nMT bundles in phase II, and shorter, yet stable, Q-nMT bundles in phase III (Fig. 2C and Sup. Fig. 2C). Moreover, “*Tub1-only*” Q-nMT bundles were thinner than WT Q-nMT bundles (Fig. 2D). Taken together, our data indicate that Tub3 is involved in MT elongation in phase II, since in the absence of this isotype, phase I MT bundles could assemble and MT bundles could get stabilized in phase III, but the second wave of MT nucleation and elongation was impaired as testified by thinner Q-nMT bundles in both phase II and III.

### Kinetochores are critical for phase I but dispensable for the maintenance of Q-nMT bundles

Kinetochore components are enriched at the Q-nMT bundle +end ((Laporte et al., 2013) and Sup Fig. 3A). We therefore tested whether kinetochore-MT attachment might play a role in Q-nMT bundle formation. First, we disrupted kinetochore-MT attachment upon glucose exhaustion using the well-established *ndc80-1* allele (Cheeseman et al., 2006; DeLuca et al., 2018; Wigge et al., 1998). As shown in Fig. 3A, only very short MT structures were detected in *ndc80-1* cells transferred to 37°C at the onset of quiescence entry, even after 4 days at non-permissive temperature (for control see Sup. Fig. 3B). Second, we focused on the chromosome passenger complex (CPC: Bir1/Survivin, Sli15/INCENP, Nbl1/Borealin and Ipl1/Aurora B), a complex known to regulate kinetochore-MT attachment dynamics (Cairo and Lacefield, 2020). We found that inactivation of *IPL1* upon entry into quiescence using the thermosensitive allele *ipl1-1* prevented Q-nMT bundle formation (Sup. Fig. 3C). The absence of Q-nMT bundles in cells harboring the NA-PP1 kinase sensitive allele *ipl1-5as* (Nerusheva et al., 2014) indicated that the Ipl1 kinase activity was required for this process (Fig. 3B and Sup. Fig. 3D-E for controls). While we could not detect Ipl1 and Nbl1 in quiescent cells, we found that Sli15-GFP and Bir1-GFP localized along the Q-nMT bundle with an enrichment at the Q-nMT bundle +end (Fig. 3C). In 4-day-old *bir1Δ* cells, Q-nMT bundle were shorter and not fully stabilized (Fig. 3D). In addition, inactivation of *SLI15* upon glucose exhaustion using the thermosensitive allele *sli15-3* (Kim et al., 1999) drastically impaired Q-nMT bundle formation (Sup. Fig. 3F). Finally, we examined the involvement of Bim1. Bim1 is a MT +end binding protein that plays a role in kinetochore MT-end on attachment (Dudziak et al., 2021; Thomas et al., 2016) and localizes along the entire Q-nMT bundle (Laporte et al., 2013). In fact, we found that that the amount of Bim1 correlated with tubulin incorporation during Q-nMT bundle formation (Sup. Fig. 3G). *BIM1* deletion alone has no effect on Q-nMT bundle formation even in the presence of Noc ((Laporte et al., 2013) and Fig. 3D). However, *bim1Δ bir1Δ* cells barely assembled MT structures, and these structures were sensitive to Noc (Fig. 3D). This demonstrates that in the absence of Bim1, Bir1 is critical for Q-nMT bundle stabilization in phase I. Of note, we found that the viability of *bim1Δ bir1Δ* cells in quiescence was drastically reduced (Sup Fig. 3H). Together, these experiments show that kinetochore-MT attachments are critical for the initiation of Q-nMT bundle formation upon entry into quiescence.

**Figure 3:**
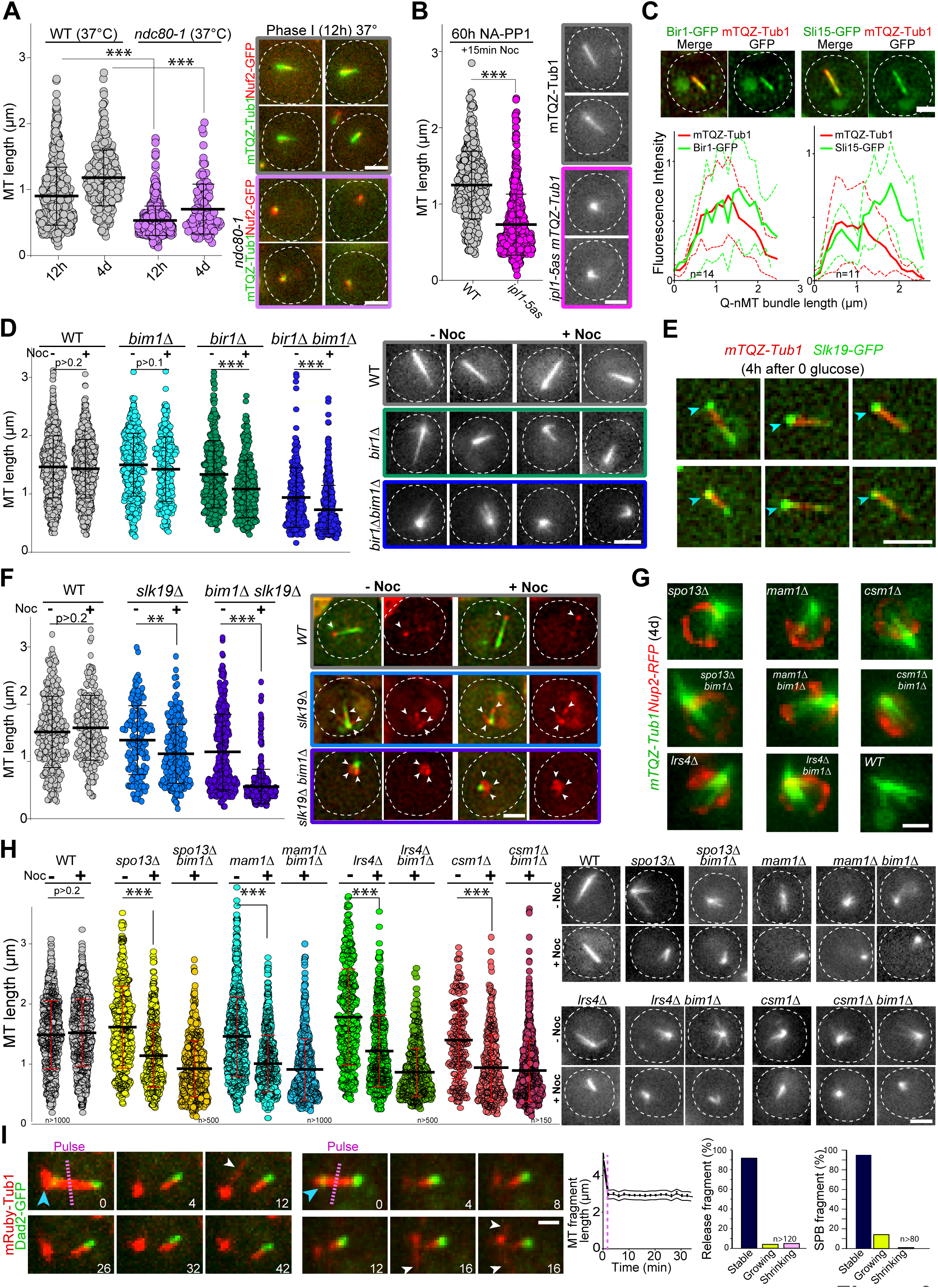
Kinetochore-kinetochore interactions are required for Q-nMT bundle formation. **(A)** Nuclear MT length distribution in WT (grey) and *ndc80-1* (violet) cells expressing mTQZ-Tub1 (green) and Nuf2-GFP (red), transferred to 37 °C upon glucose exhaustion and maintained at 37 °C for the indicated time. Cells were imaged after a 20 min Noc treatment (30 µg/ml). See Sup. Fig. 3B for the control without Noc. Each circle corresponds to the length of an individual MT structure, mean and SD are shown. A student test (t-test with two independent samples) was used to compare (**+**) or (**-**) Noc samples from the same experiment (n > 250, N = 2). The indicated p-values are the highest p-values calculated among the 2 repeated experiments. ***: p-value < 1.10^-5^. Images of representative cells incubated 12 h at 37 °C are shown, bar is 2 µm. **(B)** WT (grey) and *ipl1-5as* (pink) cells expressing mTQZ-Tub1 were treated for 60 h with 50 µM NA-PP1 after glucose exhaustion, and imaged after a 15 min Noc treatment (30 µg/ml), see Sup. Fig. 3D for control without Noc. Same statistical representation as in (A); N = 2, n > 250. Images of representative cells are shown, bar is 2 µm. **(C)** WT cells (2 d) expressing mTQZ-Tub1 (red) and Bir1-GFP or Sli15-GFP (green) were imaged. Graphs show Bir1-GFP or Sli15-GFP fluorescence intensity along normalized Q-nMT bundles (see material and method section, plain and dash lines: mean and SD respectively). Images of representative cells are shown, bar is 2 µm. **(D)** Nuclear MT length distribution in cells of the indicated genotype (4 d) expressing mTQZ-Tub1 treated or not with Noc (30 µg/ml). Same statistical representation as in (A); N = 3, n > 200. Images of representative cells are shown, bar is 2 µm. **(E)** WT cells expressing mTQZ-Tub1 (red) and Slk19-GFP (green) 4 h after glucose exhaustion. Blue arrowhead: SPB, bar is 2 µm. **(F)** Nuclear MT length distribution in cells of the indicated genotype (4 d) expressing mTQZ-Tub1 (green) and Nuf2-GFP (red) imaged before or after Noc treatment (30 µg/ml). Same statistical representation as in (A); N = 2, n > 200. Images of representative cells are shown. White arrowheads point to Nuf2-GFP dots, bar is 2 µm. **(G)** Cells of the indicated genotype (4 d) expressing mTQZ-Tub1 (green) and Nup2-RFP (red), bar is 1 µm. **(H)** Nuclear MT length distribution in cells of the indicated genotype (4 d) expressing mTQZ-Tub1 treated or not with Noc (30 µg/ml). Same statistical representation as in (A); N ≥ 2, n > 200. Images of representative cells are shown, bar is 2 µm. **(I)** Length variation of nuclear MT bundle fragments after laser ablation (pink dash line) in cells expressing mRuby-TUB1 (red) and Dad2-GFP (green). Time is in min. Images of representative cells are shown, blue arrowhead: SPB, white arrowhead: cMT. Graph indicates the variation of length in released fragments (n > 120) and histograms show the % of released or SBP attached fragments that are either stable, or that shorten or grow within a 30 min period after the laser-induced breakage. Bar is 1 µm

We then hypothesized that inter-kinetochore interactions could stabilize parallel MTs embedded in the Q-nMT bundle. To test this idea, we analyzed MT organization in s*lk19Δ* cells which are defective in kinetochore clustering (Richmond et al., 2013). In quiescence, Slk19-GFP was enriched at both ends of the Q-nMT bundles (Fig. 3E). s*lk19Δ* cells displayed shorter and thinner MT structures that were sensitive to Noc (Fig. 3F). Since Bim1 has been implicated in kinetochore-kinetochore interaction, we combined s*lk19Δ* with *bim1Δ.* MT structures detected in s*lk19Δ bim1Δ* cells barely reached ≈1 µm and disappeared upon Noc treatment. In addition, in s*lk19Δ bim1Δ* cells, kinetochores localized as a rosette around the SPB (Fig. 3F) and in s*lk19Δ bim1Δ* cell viability was severely impaired (Sup. Fig. 3H). This demonstrates that in the absence of Bim1, Slk19 is required for MT bundling and stabilization in phase I. We then focused on monopolin (Mam1, Lrs4, Hrr25 and Csm1), a complex that is known to cross-link kinetochores (Corbett et al., 2010)). These proteins were found to be expressed in quiescent cells (Sup. Fig. 3I) and when we analyzed the phenotype of *mam1Δ, lrs4Δ* and *csm1Δ* cells in quiescence, most of them displayed short MT structures often arranged in a star-like array (Fig. 3G-H and Sup. Fig. 3J). As for *bir1Δ* and *slk19Δ*, the combination of *bim1Δ* with monopolin deletion worsened MT structure length and stability (Fig. 3H) as well as cell viability in quiescence (Sup. Fig. 3H). Furthermore, cells deleted for *SPO13*, a gene encoding a protein involved in monopolin recruitment to kinetochores (Lee et al., 2004) showed similar phenotypes (Fig. 3G-H and Sup. Fig. 3H). Collectively, these experiments demonstrate that inter-kinetochore interactions are critical for Q-nMT bundle assembly.

Finally, we investigated whether kinetochore-MT interactions were required for the stability of Q-nMT bundle once it is formed. In 5-day-old WT cells, we used a UV-pulsed-laser to break the Q-nMT bundle into two pieces. In most cases, the length of both the released fragment and the fragment attached to the SPB remained constant, with the presence of dynamic cMTs after laser pulse testifying for cell viability (Fig. 3I, Sup. Fig. 3K). In contrast, as expected from previous studies (Khodjakov et al., 2004; Zareiesfandabadi and Elting, 2022), in proliferating cells, dynamic anaphase spindles promptly disassembled after the laser-induced breakage (Sup. Fig. 3L). This indicates that once Q-nMT bundles are formed, they do not require MT-kinetochore interaction to be maintained. Accordingly, inactivation of Ndc80 (Sup. Fig. 3B) or Ipl1 (Sup. Fig. 3M) in cells that were already in quiescence had no effect on the Q-nMT bundle maintenance. Thus, once formed, Q-nMT stability is established and maintained throughout its length.

### Each phase of Q-nMT bundle formation requires specific kinesins

To further dissect the molecular mechanism of Q-nMT bundle formation, we focused on kinesins. Kar3 is a kinesin that, in complex with its regulator Cik1, can generate parallel MT bundles from a MT organizing center (MTOC) both *in vitro* and in proliferative cells (Manning et al., 1999; Mieck et al., 2015; Molodtsov et al., 2016). In quiescence, we found that Kar3-3GFP localized to the SPB, but also as dots along the Q-nMT bundle (Fig. 4A). In 4-day-old *kar3Δ* cells, the majority of the detected MT structures were extremely short compared to WT (Fig. 4B and for control without Noc see Sup Fig. 4A). Similar results were obtained in *cik1Δ* but not in *vik1Δ* cells, which lack the alternative Kar3 regulators (Fig. 4B). Thus, the Kar3/Cik1 complex is required for phase I.

**Figure 4:**
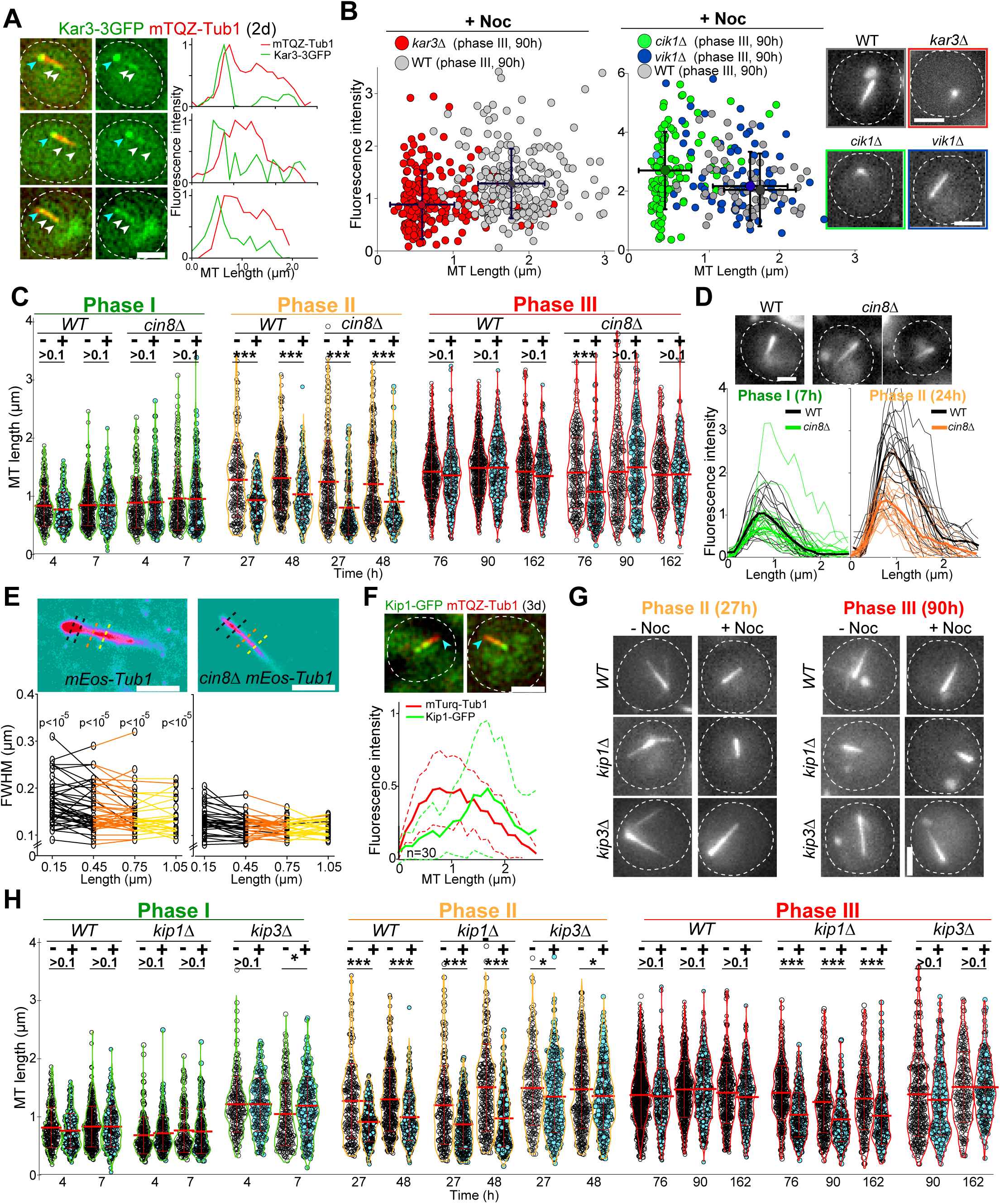
Each phase of Q-nMT formation requires a specific kinesin. **(A)** Images of representative WT cells (2 d) expressing Kar3-3GFP (green) and mTQZ-Tub1 (red) and corresponding fluorescence intensity along normalized Q-nMT bundles (see material and method section), bar is 2 µm. **(B)** Morphometric Q-nMT bundle properties distribution in 4 d WT (grey), *kar3Δ* (red), *vik1Δ* (blue), and *cik1Δ* cells (green) expressing mTQZ-Tub1 after Noc treatment (30 µg/ml) – see Sup. Fig. 4A for the control without Noc. Each circle corresponds to an individual Q-nMT bundle. Black crosses are mean and SD. Images of representative cells are shown, bar is 2 µm. **(C)** Nuclear MT length distribution in WT and *cin8Δ* cells expressing mTQZ-Tub1 treated (**+**) or not (**-**) with 15 min Noc (30 µg/ml). Each circle corresponds to the length of an individual MT structure, mean and SD are shown. A student test (t-test with two independent samples) was used to compare (**+**) or (**-**) Noc samples from the same experiment (n > 250, N = 2). The indicated p-values are the highest p-values calculated among the 2 repeated experiments. ***: p-value < 1.10^-5^. **(D)** Fluorescence intensity along Q-nMT bundles in WT and *cin8Δ* cells expressing mTQZ-Tub1 7 h and 24 h after glucose exhaustion. Thin line: intensity from an individual cell; bold line: mean intensity (n > 60 / phase). Images of representative cells are shown. Bar is 2 µm. **(E)** WT and *cin8Δ* cells expressing mEOS3.2-Tub1 were imaged using PALM Images in pseudo-colors of representative cells are shown. Full width at half maximum (FWHM) was measured at the indicated distance from the SPB. Each line in the bottom graphs corresponds to a single cell; p-value between WT and *cin8Δ* are indicated (unpaired T-test). Bar is 1 µm. **(F)** Images of representative WT cells (3 d) expressing Kip1-GFP (green) and mTQZ-Tub1 (red). Graphs show fluorescence intensity along normalized Q-nMT bundles (plain and dash lines: mean and SD respectively). Bar is 2 µm, blue arrowhead: SPB. **(G)** Representative images of WT, *kip1Δ* and *kip3Δ* cells expressing mTQZ-Tub1 treated (**+**) or not (**-**) with 15 min Noc (30 µg/ml), Bar is 2 µm. **(H)** Nuclear MT length distribution in WT, *kip1Δ* and *kip3Δ* cells expressing mTQZ-Tub1 treated (**+**) or not (**-**) 15 min with Noc (30µg/ml). Legend is the same as in (C); n > 250, N = 2; *: p-value < 1.10^-3^; ***: p-value < 1.10^-6^. MT mean length and SD are indicated.

Cin8 is a kinesin-5 that cross-links MTs (Bodrug et al., 2020; Pandey et al., 2021; Singh et al., 2018), and as such, could play a role in stabilizing Q-nMT bundles. Cin8 was barely detectable in quiescent cells (Sup. Fig. 4B) and *cin8Δ* cells assembled Q-nMT bundles that were thinner than in WT cells (Fig. 4C-E). However, these thinner bundles were stabilized in phase III (Fig. 4C and Sup. Fig. 4C-D). Taken together, these results strongly suggest that Cin8 is required for MT nucleation and/or elongation in phase II but not for MT bundle stabilization in phase III.

Yeast have an additional kinesin-5 called Kip1 (Fridman et al., 2013). In quiescence, most of the Kip1-GFP signal was observed at the Q-nMT bundle +end (Fig. 4F). In *kip1Δ* cells, phase I was slightly delayed. In phase III, while as thick as WT Q-nMT bundles, MT bundles detected in *kip1Δ* cells were not fully stabilized (Fig. 4G-H and Sup. Fig. 4E). These data demonstrate that Kip1 is required for Q-nMT bundle stabilization in phase III. Of note, in phase I and II, Q-nMT bundles were slightly longer in cells deleted for the kinesin-8 Kip3, a MT depolymerase ((Fukuda et al., 2014), see Fig. 4H) but have a WT morphology in cells deleted for Kip2, a kinesin that stabilizes cytoplasmic MTs ((Hibbel et al., 2015) and Sup. Fig. 4F). Taken together, our results demonstrate that each phase of Q-nMT bundle assembly involves specific kinesins.

### SPB duplication/separation requires Q-nMT bundle disassembly

Finally, we questioned the molecular mechanism of Q-nMT bundle disassembly upon exit from quiescence. We first found that cycloheximide prevented Q-nMT bundle disassembly indicating that this process requires *de novo* protein synthesis (Fig. 5A). Then, we measured Q-nMT bundle thickness upon depolymerization, and found that while Q-nMT bundles shortened, they did not become thinner (Fig. 5B). In agreement, Nuf2-GFP clusters found at the Q-nMT bundle +end moved back to the SPB while the Nuf2-GFP clusters localized along the Q-nMT bundle remained immobile until they are reached by the +end-associated clusters (Sup. Fig. 5A). This shows that not all MT +ends started depolymerizing at the same time, and that longer MTs depolymerized first. This finding is consistent with our model in which MTs are cross-linked along the entire length of the Q-nMT bundle.

**Figure 5:**
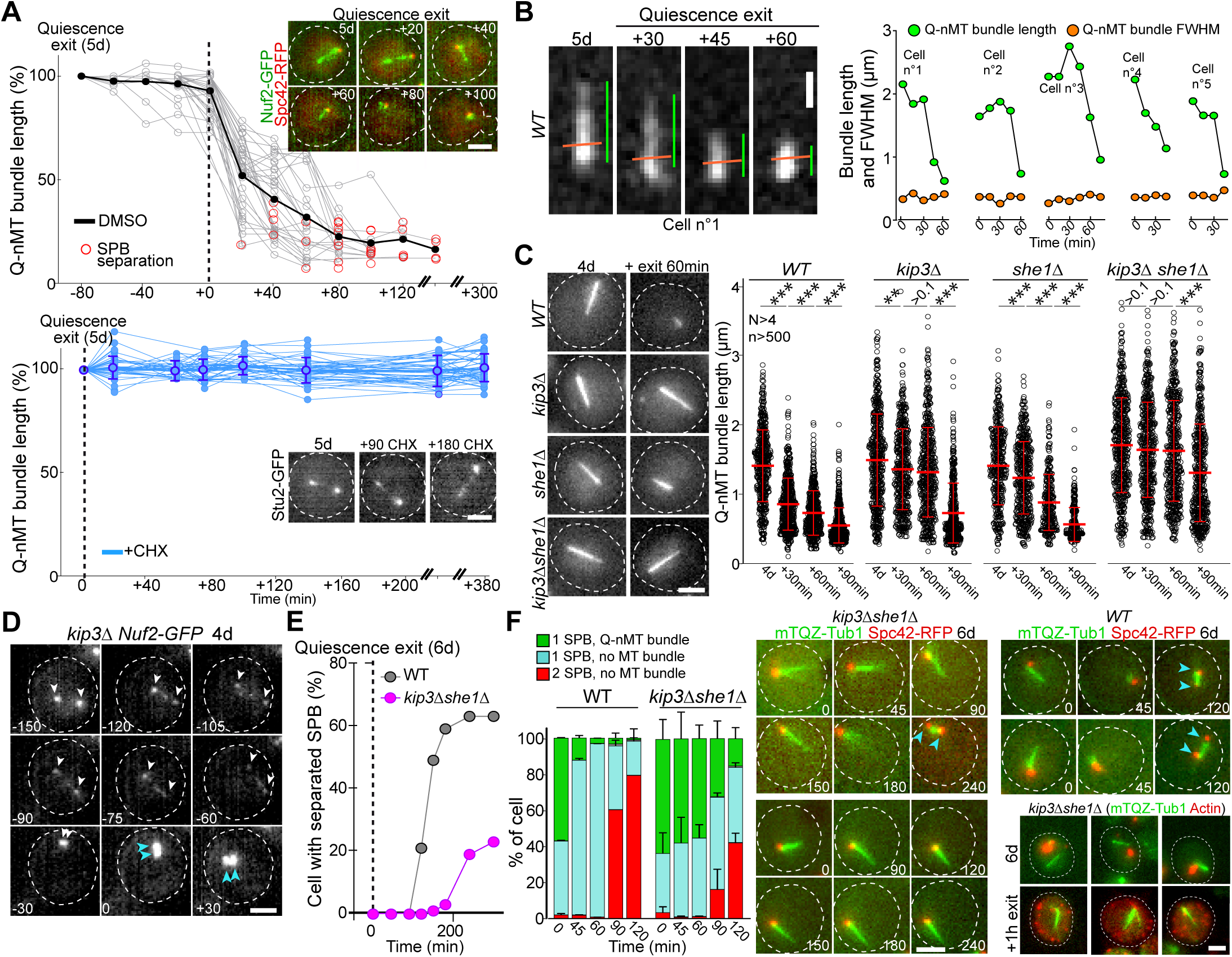
Q-nMT bundle disassembly always occurs before SPB separation upon quiescence exit. **(A)** WT cells expressing Spc42-RFP (red) (5 d) were re-fed on an YPDA microscope pad. Individual Q-nMT bundles were measured in cells expressing Nuf2-GFP treated DMSO (top panel, n=33) or in cells expressing Stu2-GFP treated with CHX (bottom panel, n=49). Each line corresponds to an individual cell. In the upper panel, time was set to zero at the onset of MT bundle depolymerisation. In the lower panel, time was set to zero when cells were deposited on the agarose pad. Images of representative cells are shown, bar is 2 µm **(B)** Q-nMT bundle length (green) and fluorescence intensity at full width half maximum (FWHM - orange) were measured upon quiescence exit in WT cells (5 d) expressing mTQZ-Tub1. Representative example of shrinking Q-nMT bundle is shown on the left, bar is 1 µm. **(C)** Cells of the indicated genotype expressing mTQZ-Tub1 were grown for 4 d, and re-fed. Q-nMT bundle length was measured at the indicated time points, 15 min after a Noc treatment (30 µg/ml), to remove dynamic cMTs that assemble upon quiescence exit. Each circle corresponds to a single cell. MT mean length and SD are indicated. A student test (t-test with two independent samples) was used to compare samples from the same experiment (n > 250, N = 4). The indicated p-values are the highest p-value calculated among the 4 repeated experiments. ***: p-value < 1.10^-5^. MT mean length and SD are indicated. Images of representative 4-day-old cells of the indicated genotype expressing mTQZ-Tub1 before and 60 min after quiescence exit are shown, bar is 2 µm. **(D)** Representative images of a *kip3Δ* cell expressing Nuf2-GFP upon quiescence exit. Blue and white arrowheads: SPBs and Q-nMT bundle extremities respectively, bar is 2 µm. **(E)** Percentage of 6-day-old WT or *kip3Δshe1Δ* cells expressing Spc42-mRFP1 with separated SPB as a function of time upon quiescence exit (n > 200). **(F)** WT and *kip3Δshe1Δ* cells (6 d) expressing Spc42-mRFP1 (red) and mTQZ-Tub1 (green) were re-fed on a YPDA microscope pad. Percentage of cells with a single SBP with or without Q-nMT bundle or with duplicated SPBs were scored (N = 4, n > 200), SD are indicated. Images of representative cells are shown, bar is 2 µm. Right bottom panel: actin (phalloidin staining, red) in *kip3Δ she1Δ* cells (6 d) expressing mTQZ-Tub1 (green) before and 1 h after quiescence exit.

Since Kip3 is the only well-characterized yeast MT depolymerase, we examined Q-nMT bundle disassembly in *kip3Δ* cells and found that it was drastically delayed (Fig. 5C-D and Sup. Fig. 5B), yet the overall depolymerization rate remained unaffected (Sup. Fig. 5B, left panel). In fact, we found that in WT cells, Kip3 needed to be resynthesized upon exit from quiescence (Sup. Fig. 5C). We searched for proteins that might be involved in Q-nMT bundle disassembly together with Kip3. Among proteins required for mitotic spindle disassembly, we found that inactivation of *CDC14*, *CDC15*, *CDH1*, *DCC1* and *TOR1* had no effect on Q-nMT bundle disassembly, as did the inactivation of Ipl1 or proteins involved in kinetochore-MT attachment (Sup. Fig. 5D-G). The only additional actor we found was She1, a dynein regulator (Bergman et al., 2012), whose deletion strongly exacerbated the *kip3Δ* phenotype (Fig. 5C and Sup. Fig. 5H). Thus, Q-nMT bundle disassembly does not rely on the canonical mitotic spindle disassembly machinery, but rather, specifically requires Kip3 and She1.

Importantly, when we followed quiescence exit at the single cell level, depolymerization of the Q-nMT bundle always preceded SPB duplication/separation (Fig. 5A) in both WT and in *kip3Δ* cells (Sup. Fig. 5B right panel). When we severely impeded Q-nMT bundle disassembly using *kip3Δ she1Δ*, we found that SPB separation was greatly delayed (Fig. 5E-F, Sup. Fig. 5I-K). Besides, the proliferation rate of *kip3Δ she1Δ* cells was the same as WT cells (Sup. Fig. 5I), and, upon quiescence exit, these mutant cells disassembled actin bodies as fast as WT cells (Sup. Fig. 5K), demonstrating that *kip3Δ she1Δ* cells were not impaired in sensing quiescence exit signals. These observations suggest that upon exit from quiescence, the Q-nMT bundle has to be disassembled prior to SPB duplication/separation.

## Discussion

Our results shed light on the precise temporal sequences that generate a Q-nMT bundle upon quiescence establishment in yeast. From bacteria to human stem cells, changes in cell physicochemical properties are known to accompany the establishment of quiescence. Among these modifications, cellular volume reduction increases molecular crowding (Joyner et al., 2016) and pH acidification both increases viscosity and causes changes in macromolecule surface charges (Charruyer and Ghadially, 2018; Jacquel et al., 2021; Munder et al., 2016; Persson et al., 2020). Much evidence suggests that these modifications act as triggers for the auto-assembly of several types of enzyme-containing granules (Munder et al., 2016; Petrovska et al., 2014; Rabouille and Alberti, 2017), or complex structures such as P-bodies and Proteasome Storage Granules (Peters et al., 2013; Jacquel et al., 2021; Currie et al., 2023). Recently, Molines and colleagues demonstrated that cytoplasmic viscosity modulates MT dynamics *in vivo* (Molines et al., 2022). Here, we showed that the formation of the Q-nMT bundle does not depend on changes in the physicochemical properties that cells experience at the onset of quiescence establishment (Sup. Fig. 1). In fact, dynamic cMTs and a stable Q-nMT bundle can be observed simultaneously in a quiescent cell (Sup. Fig. 1I), while physicochemical properties evolve in parallel in both the nucleus and the cytoplasm (Joyner et al., 2016). Importantly, cellular volume reduction, pH acidification and increased viscosity are observed within minutes upon glucose starvation (Joyner et al., 2016) whereas the Q-nMT bundles require a couple of days to be fully assembled (Fig. 1). Furthermore, Q-nMT bundle formation cannot be delayed in time (Fig. 1H), and thus likely depends on a transient signal emitted upon glucose depletion. We speculate that the SPB may act as a platform that integrates this signal, and transfers it to the MT machinery in order to assemble a stable MT structure.

As it is required for the onset of Q-nMT bundle formation (Fig. 3B), the CPC component Aurora B/Ilp1, a kinase that plays a key role at the MT-kinetochore interface in response to the tension status, could be one of the targets of the aforementioned nutritional signal. In parallel, upon quiescence establishment, chromosome hyper-condensation (Guidi et al., 2015; Rutledge et al., 2015) could modify the tension at the kinetochore/MT interface and as such could contribute to the initiation of phase I (Fig. 6b). Besides, Bim1 has also been implicated in MT-kinetochore attachment (Dudziak et al., 2021; Akhmanova and Steinmetz, 2015). Although its deletion alone has no effect on Q-nMT bundle formation, its role becomes critical for phase I when kinetochore-MT interactions are already destabilized by the absence of other CPC components, other than Ipl1, such as Bir1 (Fig. 3D). Taken together, these observations suggest that kinetochore-MT interactions are essential for phase I initiation, as confirmed by the absence of Q-nMT bundle in the *ndc80-1* mutant (Fig. 3A).

**Figure 6:**
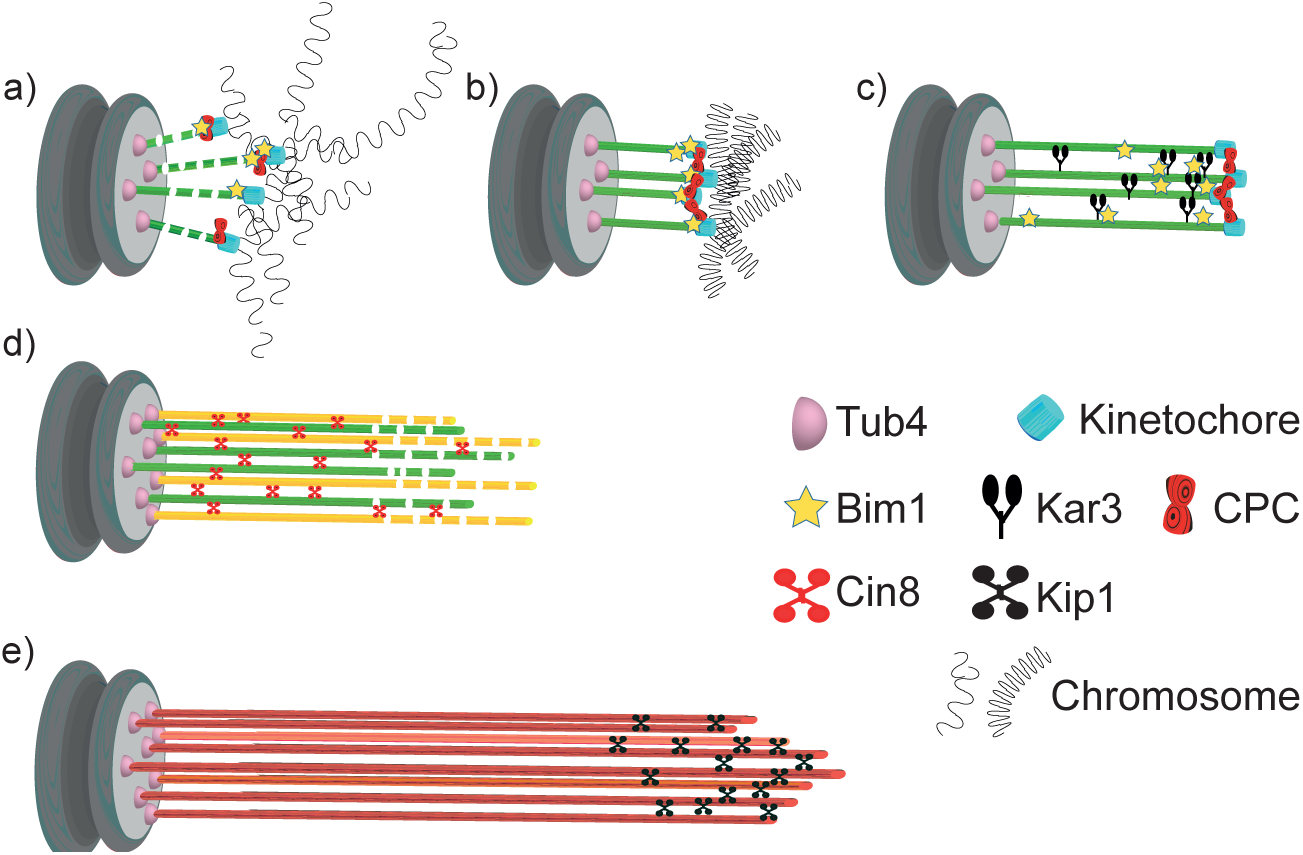
Model for Q-nMT bundle assembly. (a) In G1, the nucleus is in a Rabl-like configuration. (b) Upon quiescence establishment, chromosomes get condensed. MT-kinetochore interaction and Ilp1 are required for the onset of phase I. (c) Kar3 and its regulator Cik1 are essential to initiate Q-nMT bundle elongation. Although deletion of *BIM1* has no effect, it becomes critical for phase I if kinetochore-MT interactions are destabilized by the absence of Chromosome Passenger Complex components. Kinetochore clustering by the monopolin complex and Slk19 is needed to maintain MT bundling while phase I-MTs elongate. During phase I, Tub4 accumulates at the SPB. (d) In phase II, a second wave of MT nucleation and elongation occurs. Phase II-MTs are concurrently stabilized along pre-existing phase I-MTs, in a Cin8-dependent manner. Phase I and phase II MTs +ends (>1 µm) remain dynamic until the full-length Q-nMT bundle stabilization is reached via the action of Kip1, about 2 days after glucose exhaustion (e).

In phase I, SPB-anchored MTs elongate and are concomitantly stabilized (Fig. 1A-C). This step requires the kinesin-14 Kar3 and its regulator Cik1 (Fig. 4B). During mating (Molodtsov et al., 2016), and in early mitosis, when half-spindles form (Kornakov et al., 2020), Kar3/Cik1, together with Bim1, align and cross-link growing MTs along existing MTs, thereby promoting the organization of MTs into parallel bundles. In mitosis however, the kinesin-14/EB1 complex promotes MT dynamics (Kornakov et al., 2020). It remains to be determined whether the robust MT stabilization in phase I, depends on the modification of the kinesin-14/EB1 complex properties and/or on additional specific MT cross-linker(s) (Fig. 6c).

In quiescence, deletion of either monopolin or *SLK19* results in the formation of short, shattered and flared MT structures, phenotypes that are exacerbated by the deletion of *BIM1* (Fig. 3F-H). Since monopolin and Slk19 are involved in kinetochore clustering (Plowman et al., 2019; Rabitsch et al., 2003; Tóth et al., 2000; Lee et al., 2004; Mittal et al., 2019; Movshovich et al., 2008; Richmond et al., 2013; Zeng et al., 1999), we speculate that kinetochore-kinetochore interaction may not only help prevent depolymerization of individual MTs, but also constrain elongating phase I-MTs and facilitate their concomitant cross-linking (Fig. 6d).

With the initiation of phase I, Tub4 begins to accumulate at the SPB (Fig. 1E and Sup. Fig. 1F) to enable the second wave of MT nucleation observed in phase II. As phase II-MTs elongate, they are simultaneously stabilized along pre-existing phase I-MTs (Fig. 6d). This step relies on the kinesin-5/Cin8 as Q-nMT bundles assembled in *cin8Δ* cells are thinner than WT Q-nMT bundles (Fig. 4C-E). Intriguingly, *in vitro,* the monomeric form of the vertebrate kinesin-5, Eg5, promotes MT nucleation and stabilizes lateral tubulin-tubulin contacts (Chen et al 2019). We assume that in the absence of Cin8, either nucleation of phase II-MTs cannot occur, or phase II-MTs can elongate, but cannot be cross-linked and stabilized to pre-existing phase I-MTs as they grow, and thus rapidly depolymerize. Absence of bundle thickening in phase II is also observed in the “*Tub1-only*” mutant *i.e.* in the absence of Tub3 (Fig. 2D). Tubulin isotypes or tubulin PTMs are known to promote the recruitment of specific kinesins and thereby modify MT properties, see for example (Sirajuddin et al., 2014; Peris et al., 2009; Dunn et al., 2008). Could Tub3 be important for Cin8 recruitment or function during phase II?

Following phase II, MTs elongate, but their distal parts (>1 µm) are not yet stable (Fig. 1A). Stabilization of full-length Q-nMT bundles is achieved approximately 4 days after glucose exhaustion. This step depends on the kinesin-5/Kip1 (Fig. 4G-H and Sup. Fig. 4E) which is essential for cross-linking and stabilizing elongating MTs during phase III (Fig. 6e). Accordingly, we observed a Kip1 enrichment at the Q-nMT bundle +ends (Fig. 4F). Thus, consistent with their ability to form homo-tetramers capable of cross-linking (Acar et al., 2013; Kapitein et al., 2005; Scholey et al., 2014; Weinger et al., 2011) and stabilizing parallel MTs (Kapitein et al., 2005; Shimamoto et al., 2015; Yukawa et al., 2020), and in addition to their role of sliding anti-parallel MTs, the two yeast kinesin 5 are essential for Q-nMT bundle formation. However, in quiescence, as well as in mitosis, Kip1 and Cin8 have non-equivalent functions (Roostalu et al., 2011; Shapira and Gheber, 2016).

Thus far, our results indicate that the mechanism of Q-nMT bundle formation is distinct from that of mitotic spindle assembly. Similarly, we show that Q-nMT bundle disassembly does not involve the same pathway as mitotic spindle disassembly, as it requires Kip3 (Fig. 5C) but is independent of the anaphase-promoting complex and Aurora B/Ipl1 (Sup. Fig. 5D-G). Interestingly, the longest MTs begin to depolymerize from their +ends first. Then, when they reach the +ends of shorter MTs, the shorter ones proceed to depolymerize alongside them (Fig. 5B and Sup. Fig. 5A). This cooperative behavior has been observed *in vitro* within parallel MT bundles (Laan et al., 2008). Our data demonstrate that such a comportment does exist *in vivo*.

Finally, we show that the Q-nMT bundle is required for the survival of WT cells in quiescence (Sup. Fig. 3H). While we do not yet know why this structure is important for chronological aging, it is clear that the presence of the Q-nMT bundle modifies the organization of the nucleus (Laporte et al., 2013; Laporte and Sagot, 2014; Laporte et al., 2016). Since chromatin organization influences gene expression, one hypothesis could be that the presence of the Q-nMT bundle is required for the expression of genes necessary for cell survival in quiescence. Another attractive hypothesis is that the Q-nMT bundle could be the yeast analog of the mammalian primary cilium. Indeed, these two structures are formed upon proliferation cessation, they are both composed of highly stable parallel MTs, their formation involves MT motors, and they are both templated from the centriole/SPB. More importantly, we show here that Q-nMT bundle disassembly always occurs prior to SPB separation upon exit from quiescence (Fig. 5E-F and Sup. Fig. 5J). Thus, the Q-nMT bundle disassembly may control reentry into the proliferation cycle, just as the primary cilium does in ciliated mammalian cells (Goto et al., 2017; Kim and Tsiokas, 2011).

## Materiel and Methods

### Yeast strains, plasmids and growth conditions

All the strains used in this study are isogenic to BY4741 (*mat a, leu2Δ0, his3Δ0, ura3Δ0, met15Δ0)* or BY4742 (*mat alpha, leu2Δ0, his3Δ0, ura3Δ0, lys2Δ0),* available from GE Healthcare Dharmacon Inc. (UK), except for Fig. 2, in which strains of the S288c background were used (Nsamba et al., 2021) and Sup Fig 1F in which the W303 background was used. BY strains carrying GFP fusions were obtained from ThermoFisher Scientific (Waltham, MA, USA). Integrative plasmids p*Tub1-mTurquoise2-Tub1*, p*Tub1-wasabi-Tub1*, p*Tub1-mRUBY2-Tub1*, p*Tub1-mEOS2-Tub1* were a generous gift from Wei-Lih Lee (Markus et al., 2015). The strain expressing SPC42-mRFP1 is a generous gift from E. O’Shea (Huh et al., 2003). The strain expressing TUB4-mTQZ is a generous gift from S. Jaspersen (Burns et al., 2015). The *ipl1-5as* is a generous gift from A. Marston (Nerusheva et al, 2014). *Ipl1-1*, *ndc80-1* and *sli15-3* are generous gift form C. Boone. Three copies of eGFP in tandem were integrated at the 3’ end of *DAD2*, *TUB3* or *KAR3* endogenous loci respectively. Plasmids for expressing Nup2-RFP (p3695) or Bim1-3xGFP (p4587) from the endogenous locus were described in (Laporte et al., 2013).

For Fig. 1F and Sup. Fig. 1H, *pHIS3:mTurquoise2-Tub1+3’UTR::LEU2* was integrated at *TUB1* locus, then a pRS303-*ADH2p-mRuby2-Tub1* was integrated at the *HIS3* locus. To generate this plasmid, the *ADH2* promoter was amplified from yeast genomic DNA flanked with NotI and SpeI restriction sites and inserted in pRS303. The mRuby2-Tub1+3’UTR, including the 116-nucleotide intron and 618 nucleotides downstream the stop codon was cloned between the SpeI and SalI sites.

Yeast cells were grown in liquid YPDA medium at 30 °C in flasks as described previously in (Sagot et al., 2006) except for experiments using thermosensitive strains, where cells were first grown at 25 °C then shifted at 37 °C for indicated time before imaging.

Experiment in Sup. Fig. 1J was performed as described in (Orij et al., 2009, Peters et al., 2013). In brief, proliferating cells at an OD_600nm_ of 0.5 were transferred in Hepes buffer (25 mM Hepes, pH 7.4, 200 mM KCl, 1 mM CaCl_2_, and 2 % dextrose), buffered either at pH 4 or 7.5, in the presence of 100 µM CCCP (Sigma-Aldrich). After 150 min at 30 °C with shaking, cells were imaged (for mTQZ-Tub1) or fixed with formaldehyde and stained using Alexa Fluor 568– phalloidin (Invitrogen) as described (Sagot et al., 2006).

For live cell imaging, 2 μL of the cell culture were spotted onto a glass slide and immediately imaged at room temperature.

For quiescence exit (Fig. 5), cells were first incubated 2-3 min in liquid YPD and then 2 µL were spread onto a 2 % agarose microscope pad containing YPD. Individual cells were imaged every hour, up to 6 or 12 h at 21°C. For quiescence exit in presence of cycloheximide (Fig. 5A), cells were pre-incubated 30 min in the presence of the drug prior quiescence exit.

Immunofluorescence was done as in (Laporte et al. 2016), and AlexaFluor Phalloidin (Invitrogen) staining as in (Sagot et al., 2006).

Cycloheximide was used at 180 µM (Sigma-Aldrich), Nocodazole was used at 30 µg/mL (7.5 µM) (Sigma-Aldrich), NA-PP1 was used at 50 µM (Sigma-Aldrich).

### Fluorescence Microscopy

Cells were observed on a fully-automated Zeiss 200M inverted microscope (Carl Zeiss, Thornwood, NY, USA) equipped with an MS-2000 stage (Applied Scientific Instrumentation, Eugene, OR, USA), a Lambda LS 300 W xenon light source (Sutter, Novato, CA, USA), a 100X 1.4NA Plan-Apochromat objective, and a 5-position filter turret. For RFP and mRuby2 imaging we used a Cy3 filter (Ex: HQ535/50 nm – Em: HQ610/75 nm – BS: Q565 nm lp). For GFP imaging, we used a FITC filter (excitation, HQ487/25 nm; emission, HQ535/40 nm; beam splitter, Q505 nm lp). For mTurquoise2 imaging we used a DAPI filter (Ex: 360/40 nm – Em: 460/50 nm– BS: 400 nm). All the filters are from Chroma Technology Corp. Images were acquired using a CoolSnap HQ camera (Roper Scientific, Tucson, AZ, USA). The microscope, camera, and shutters (Uniblitz, Rochester, NY, USA) were controlled by SlideBook software 5. 0. (Intelligent Imaging Innovations, Denver, CO, USA).

For PALM (Fig. 4E) a Nikon Ti-Eclipse equipped with iLas2 TIRF arm, laser diodes (405, 488, 532, 561, 642 nm), a 100x 1.49 oil (TIRF) objective connected to a EMCCD Camera Photometrics Evolve was used.

For expansion microscopy (Sup Fig. 2A and 3A), spheroplasts were obtained as described in (Laporte et al., 2016). After washing in 10 mg/mL NaBH4 in PEMS for 10 min, spheroplasts were seeded on 12 mm round poly-L-lysine coated coverslips, washed twice for 30 min in 100 μL PEM-BAL (PEM + 1 % BSA) and incubated with the primary antibodies (anti-GFP from mouse; Roche, ref. 11814460001, 1/50; anti-alpha-tubulin from rat; YOL1/34, Abcam, ref. Ab6161, 1/100) for 1 hour at 37 °C. Cells were then washed three times in PEM-BAL and incubated with the secondary antibodies (Goat anti-mouse AlexaFluor®488 1/200 and Donkey anti-rat AlexaFluor®555; A11029 and A21434 respectively, ThermoFisher Scientific, Waltham, MA) for 45 min at 37 °C. Cells were then washed three times with PBS. Cells were processed for Expansion Microscopy as previously described in (Bahri et al., 2021). Spheroplasts were incubated for 10 min in 0.25 % GA in PBS, washed in PBS three times for 5 min, and processed for gelation. A drop of 100 µL of ExM MS (8.625 % (wt/wt) SA, 2.5 % (wt/wt) AA, 0.15 % (wt/wt) BIS, 2 M NaCl in 1× PBS) was placed on the chilled Parafilm, and coverslips were put on the drop with cells facing the solution and incubated for 3 min. Then the coverslips were transferred to 35 μL of ExM MS supplemented with 0.2 % APS and 0.2 % TEMED, with the initiator (APS) added last. Gelation proceeded for 3 min on ice, and samples were incubated at 37 °C in a humidified chamber in the dark for 1 h. Then coverslips with attached gels were transferred into a six-well plate for incubation in 2 mL of digestion buffer (1× TAE buffer, 0.5 % Triton X-100, 0.8 M guanidine hydrochloride, pH ∼8.3) supplemented with DAPI (1 µg/mL) for 10-15 min at 37 °C, until the gels detached. Fresh proteinase K at 8 units/mL was then added and samples incubated at 37 °C for 30 min. Finally, gels were removed and placed in 10 mL petri dishes filled with ddH2O for expansion. Water was exchanged at least twice every 30 min, and incubated in ddH2O overnight at RT. Gels expanded between 4× and 4.2× according to SA purity. Expanded cells were imaged with a UPlanS Apo 100X/1.4 oil immersion objective in an Olympus IX81 microscope (Olympus, Tokyo, Japan). For structured illumination microscopy (Sup Fig 3A), a ZEISS Elyra 7 Lattice SIM was used.

Laser ablation (Fig. 3I and Sup. Fig. 3K-L) was performed at room temperature with a 100X oil Plan-Apochromat objective lens (NA 1.4) and an Axio-Observed.Z1 microscope (Carl Zeiss) equipped with a spinning disk confocal (Yokogawa), an EMCCD Evolve camera (Photometrics and Roper Scientific) and 491 nm (100 mW; Cobolt calypso) and 561 nm (100 mW; Cobolt Jive) lasers. Images were acquired with Metamorph software (Roper Scientific). Every 30 to 120 s, a Z-series of 0.4 µm steps were acquired. A 355 nm microchip laser (Teem Photonics) with a 21 kHz repetition rate, 0.8 µJ energy/pulse, 2 kW of peak power and 400 ps pulse width, powered with an iLas2 PULSE system (Roper Scientific) was used between 10 to 40 % power with one pulse of a spot length of 100 points. Breakage was considered as successful if a non-alignment between the two remaining Q-n MT bundle fragments was observed.

Z-stacks were deconvolved using the Deconvolution Lab plugin (Fig. 1D, F; Fig. 3C, E, F; Fig. 4A, F; Sup. Fig. 1E; Sup. Fig. 2A and Sup. Fig. 4B).

In Fig. 1H, F, Fig. 2B-C, Fig. 3C, Fig. 4D, G, Sup. Fig. 1I, Sup. Fig. 2E and Sup. Fig. 3A, a fuzzy fluorescence signal was detected in some cell cytoplasm using the GFP long pass filter set. This signal is not GFP but rather due to a non-specific yellow background fluorescence.

### Image analysis

Distribution and associated statistics were performed using GraphPad Prism 5 (GraphPad Software, Inc. La Jolla, USA) and Excel (Microsoft). Unless specified, a student test (t-test with two independent samples) was used to compare two conditions. Among p-values obtained for biological repetitions of the same experiment (Fig. 1A; Fig. 2B-C, Fig. 3A, D, F, H; Fig. 4C, H, and Fig. 5C), the indicated p-values are the highest calculated among the repeated experiments. p-values above 0,05 are indicated, while p-values below 0,05 are indicated using ** (0,05 > p-value > 1 10^-4^) and *** if the p-values was less than 1 10^-5^). In histograms and scatter dot plots, the mean is shown and the error bars indicate SD.

MT length was measured on MAX-projected image using imageJ.

For MT fluorescence intensity measurements (Fig. 1B, Fig. 2D, Fig. 4D, Sup. Fig. 1A, and Sup. Fig. 4C, E), a line scan (i1) of 4 pixels width containing both FP signal and background was drawn along MTs on sum projection image (3-4 Z-plans) using ImageJ. A line of 8 pixels width (at the same location) was drawn in order to calculate the intensity of the surrounding background (i2). The real intensity (ir) was calculated as follow ir = [i1 - ([(i2 x i2_surface_) - (i1 x i1_surface_)] / (i2_surface_ - i1_surface_)) * i1_surface_]. We arbitrarily set the ‘zero’ at 0,5µm before the fluorescence increased onset on the SPB side in order to align all intensity measurements. Similarly, to measure Tub4 fluorescence intensity (Fig. 1E and Sup. Fig. 1F). two boxes (i1) and (i2) were drawn around the fluorescence signal. Intensity was calculated as follow: int = [i1 - ([(i2 x i2_surface_) - (i1 x i1_surface_)] / (i2_surface_ - i1_surface_)) * i1_surface_].

For morphometric Q-nMT bundle property distribution (Fig. 1C, Fig. 4B and Sup. Fig. 4A, D) the mean fluorescence intensity was measured in each individual cell, in a finite area localized adjacent to the SPB (0,4 µm – 3-4 Z stack sum projected) as an estimate of Q-nMT bundle width, and plotted as a function of the corresponding bundle length.

For Nuf2-GFP fluorescence intensity measurement at the SPB (Sup. Fig. 1E, bottom panel), GFP and RFP line scan measurements were done on a 4 Z-planes sum projection. The “SPB localization zone” was defined as the length ending 250 µm after the brightest RFP signal of Spc42-RFP. The remaining Q-nMT bundle zone was defined as the “+end zone”. The total Nuf2-GFP signal detected along the Q-nMT bundle was set to 100 % to determine the percentage of the signal measured at the “SPB localization zone” and the “+end zone”.

Normalized Q-nMT bundle length (Fig. 3C, and Fig. 4A, F) was calculated after line scan intensity measurement. mTQZ-Tub1 intensity slopes were first aligned on their inflexion points. To compare Q-nMT bundles with different lengths, we first sorted Q-nMT bundles < 1.8 µm. After an artificial isotropic expansion, we fit all MT structures to the same length.

Then, the corresponding mean intensity of Q-nMT bundles (mTZQ-Tub1) and GFP signal were calculated.

To measure Q-nMT bundle depolymerization rate (Fig. 5A and Sup. Fig. 5B), individual Q-nMT bundle lengths were measured over time. The first length measurement was set to 100 %. A fluorescence drop above or equal to 25 % defines the inflexion point of the slope, and was used to align the length measurements for Q-nMT bundles.

The FWHM (Full Width at Half Maximum, Fig. 5B and Sup. Fig. 1B) was calculated by measuring fluorescence intensity of a line crossing perpendicularly a Q-nMT bundle, at 0.5 µm from the SPB, using a sum projection image. After fitting the intensity level with Gaussian distribution and obtaining the standard deviation (σ) value, FWHM was calculated using the equation 2√(2ln2)xσ.

### Western blots

Western blots were done as described in (Sagot et al., 2006) using anti-GFP antibodies (Abcam); anti-Tat1 antibodies (a generous gift from J-P. Javerzat) and antibody against the budding yeast Act1 or Sac6, generous gifts from B. Goode and anti-TagRFP (mRuby2) antibodies, a generous gift from M. Rojo.

### Phenotypical analysis of cells without Q-nMT bundle

WT cells were grown to glucose exhaustion (OD ∼6.5) (Fig. 1F) or for 5 d (Sup. Fig. 1L), washed with “old YPDA” (YPDA medium in which cells were grown for 4 d and then filtered to remove cells) and treated with either DMSO or Noc (30µg/mL) for 24 h. Cells were then washed twice in “old YPDA” before being incubated in the same medium. Dead cells were identified using methylene blue staining. The capacity of cells to exit quiescence (Sup. Fig. 1L, right panel) was scored after micro-manipulation of live cells (cells not stained with methylene blue) as described in (Laporte et al., 2011) (N=4, n>100).

## Supplemental data

**Supplementary Figure 1** further describes the 3 steps of Q-nMT bundle formation. It shows the impact of physicochemical cell properties on MT stabilization and demonstrates that Q-nMT bundle is required for both cell viability in quiescence and quiescence exit fitness.

**Supplementary Figure 2** describes the impact of the alpha tubulin level on Q-nMT bundle assembly.

**Supplementary Figure 3** demonstrates that kinetochore-MT interactions are not required for Q-nMT bundle maintenance and that mutants affected for kinetochore-kinetochore interactions have reduced viability in quiescence.

**Supplementary Figure 4** further describes the impact of kinesin deletion on Q-nMT bundle morphometric parameters.

**Supplementary Figure 5** explores Q-nMT bundle disassembly in mutants involved in mitotic spindle dismantlement.

## Acknowledgments

The authors would like to thank the Bordeaux Imaging center for the help in super-resolution imaging. We express our gratitude to E. O’Shea, A. Marston, S. Jaspersen, J-P. Javerzat, Wei-Lih Lee, C. Boone and B. Goode for sharing reagents and M. Rojo for providing us with the mCherry specific antibodies. We thank Michaël Gué (Zeiss) for helping us a with the lattice SIM Elyra 7 microscope. We would like to thank J-P. Javerzat for helpful and constructive discussions about our work. DL, AML, JD and IS were supported by a grant from the ANR-21-CE13-0023-01, la Ligue Contre le Cancer Régionale – Dordogne grant #193366 and the CNRS. MG and EN were supported by a National Science Foundation grant number MCB-1846262. AR was supported by the Conseil Régional de Nouvelle Aquitaine (#20111301010) and the CNRS.

## Author Contributions

D. Laporte did all the experiments described in this manuscript, except Fig. 1F, 1H, Fig. 5E-F, Sup. Fig. 1H, Sup. Fig. 3E, J and Sup. Fig. 5I to K that were performed by A. Massoni-Laporte, together with all image deconvolution, and Fig. 4E and Sup. Fig. 2A and 3A that were done by J. Dompierre. A. Royou helped for pulse-laser experiments of Fig. 3I and Sup Fig. 3K and L. C. Lefranc did molecular biology and western blots. D. Mauboules performed yeast genetic for Fig. 2. M. L. Gupta Jr and E. T. Nsamba created the Tub1-only mutant (Fig. 2). L. Gal and M. Schuldiner performed high throughput deletion screen. IS and DL designed and supervised the experiments. IS and DL wrote the manuscript.

## Abbreviations

MT: microtubule
SPB: spindle pole body
GTP: guanosine triphosphate
MTOC: microtubule organizing center
ɣ-TuC: ɣ-tubulin complex
MAPs: microtubule associated proteins
Q-nMT bundle: quiescence specific nuclear microtubule bundle
CPC: chromosome passenger complex

## Supplementary Figure legends

**Supplementary Figure 1 (A)** Fluorescence intensity along MT structures in WT cells expressing mTQZ-Tub1. Mean intensity measurement for half pre-anaphase mitotic spindles (violet, n = 60), phase I (green, n = 19), and phase II (orange, n= 22) Q-nMT bundles. A line scan along the MT structure for an individual cell is shown as thin line, the mean as a bold line, all the lines being aligned at 0,5 µm before the fluorescence intensity increase onset on the SPB side.

**(B)** Individual Q-nMT bundle full width measured at half-maximum in WT cells (4 d) expressing mTQZ-Tub1 before or after 1 h after Noc treatment. A paired t-test was use to compare the two conditions (n > 40).

**(C)** Nuclear MT length in WT haploid (mat a and mat α) or diploid (mat a/mat α) cells expressing mTQZ-TUB1. Each circle represents an individual cell. Mean and SD are shown in red.

**(D)** Nuclear MT length in WT proliferating cells expressing mTQZ-Tub1 transferred to water for the indicated time and treated (**+**) or not (**-**) 15 min with 30 µg/ml Noc. Images of representative cells are shown, bar is 2 µm.

**(E)** WT cells expressing Spc42-RFP (red) and mTQZ-Tub1 (green, top panel) or Nuf2-GFP (green, bottom panel) upon entry into quiescence. The percentages indicate the relative amount of Nuf2-GFP detected at the SBP (blue arrowhead), bar is 1 µm.

**(F)** Tub4-mTQZ fluorescence intensity in WT cells of the W303 background. The mean and SD are shown; t-test was used to compare independent samples (N = 3, n > 150), ***: p-value < 1.10^-5^.

**(G)** Fluorescence intensities in WT cells (4 d) expressing Tub4-mTQZ and mRuby-Tub1. The Tub4-mTQZ signal at the SBP is plotted as a function of the mRuby signal along the Q-nMT bundle. Each circle represents an individual cell (n = 60).

**(H) Top panel:** western blot using anti-GFP and anti-RFP antibodies on total protein extracts obtained from the indicated cells upon entry into quiescence. Anti-Ade13 antibodies were used as loading control.

**Bottom panel**: images of representative WT cells expressing only pTUB1-mTQZ-Tub1 (green) or only pADH2-mRuby-Tub1 (red) upon entry into quiescence as a control of cross channel fluorescence, bar is 2 µm.

**(I)** WT cells expressing mRuby-Tub1 (4 d) displaying both dynamic cMTs (red arrowhead) and a Q-nMT bundle (time is in min, bar is 1 µm).

**(J)** An artificial fluid to solid-like phase transition in proliferating cells does not induce MT stabilization. Proliferating WT cells expressing mTQZ-Tub1 (green) and Spc42-RFP (red) with fluid (pH = 7.5) or solid-like (pH = 4) cytoplasm/nucleoplasm (see materials and method) were incubated 15 min with 30 µg/ml Noc (left) or stained with phalloidin to detect actin (right). Indicated phenotypes were quantified (left and right graphs, N = 2, n > 115, mean and SD are indicated). Bar is 2 µm.

**(K)** Cells can assemble dynamic cMTs (red arrowhead) in quiescence. 2-day-old WT cells expressing mRuby-Tub1 were imaged after a 15 min Noc pulse-chase (30 µg/ml, time is in min). Red arrowhead: cytoplasmic MT, blue arrowhead: SBP, bar is 2 µm.

**(L) Left graphs:** WT cell viability after a 24h-pulse (grey rectangle) with Noc (30 µg/ml, circles) or DMSO (triangles), followed by a chase using carbon exhausted medium, the Noc pulse being done either upon glucose exhaustion or 5 days after glucose exhaustion. Each circle/triangle is the percentage obtained for an independent experiment in which n > 200 cells were scored.

**Right graph:** live cells, as attested by a clear staining after a methylene blue treatment, were micro-separated 8 days after the Noc pulse done either upon glucose exhaustion or 5 d after glucose exhaustion, and allowed to give rise to a colony on a YPDA plate. Each circle/triangle is the percentage obtained for an independent experiment in which n > 100 cells were micro-separated. Images of representative YPDA plates after 3 d of growth at 30°C are shown.

**Supplementary Figure 2**

**(A)** Expansion microscopy images of WT cells (4 d) expressing Tub3-3GFP and immuno-stained with GFP (green) and α-tubulin (red) YOL1/2 antibodies, bar is 1 µm.

**(B)** Western blot using α-tubulin YOL1/2 antibody on total protein extracts from the indicated cells grown for 4 d. Actin was used as a loading reference. Numbers indicate the tubulin/actin ratio (mean for N = 8).

**(C)** Nuclear MT length in WT, *tub3Δ* and tub1-only cells expressing mTQZ-Tub1, treated or not with 30 µg/ml Noc. The three independent experiments used in Fig. 2B and Fig. 2C are shown. P-values for Anova or unpaired t-test are indicated, ***: p-value < 1.10^-5^.

**(D)** Nuclear MT length distribution in *yke2Δ*, *pac10Δ* or *gim3Δ* cells (4 d) and in corresponding WT cells expressing Nuf2-GFP (dark grey bars) or Bim1-3GFP (light grey bars). Right panel: western blot using an anti-α-tubulin antibody on total protein extracts from cells of the indicated genotype (4 d). Actin was used as a loading reference. The tubulin/actin ratio is indicated (mean of N=4). Bar is 2 µm.

**(E)** Nuclear MT length distribution in *pac2Δ*, *cin2Δ* or *cin4Δ* cells (4 d) and in corresponding WT cells expressing Bim1-GFP (dark grey bars). Right panel: western blot using an anti-α-tubulin antibody on total protein extracts from cells of the indicated genotype (4 d). Actin was used as a loading reference. The tubulin/actin ratio is indicated (mean for N=4). Bar is 2 µm.

**Supplementary Figure 3**

**(A) Upper panel:** WT cell (4 d) expressing Dad2-GFP and immuno-stained with GFP (green) and α-tubulin (red) YOL1/2 antibodies imaged using expansion microscopy. Blue is DAPI.

**Lower panel:** WT cell (4 d) expressing Dad2-GFP (green) and mRuby-Tub1 (red), imaged using SIM. Bar is 1 μm.

**(B)** Nuclear MT length distribution in cells of the indicated genotypes, grown for 3 d at 25 °C, shifted to 37 °C or 25° C for 48 h, and treated 15 min or not with 30 µg/ml Noc. Each circle corresponds to the length of an individual MT structure, mean and SD are shown. ***: p-value < 1.10^-6^. Images of representative cells are shown, bar is 2 µm.

**(C)** WT (grey) and *ipl1-1* cells (pink) expressing mTQZ-Tub1 were shifted upon glucose exhaustion to 37 °C for 48 h, and imaged after a 20 min Noc treatment (30 µg/ml). Same legend as in (B). Images of representative cells are shown, bar is 2 µm.

**(D)** Nuclear MT length in cells of the indicated genotypes, grown for 60 h in DMSO or 50 µM NA-PP1, treated 15 min or not with 30 µg/ml Noc. Same legend as in (B). Images of representative cells are shown, bar is 2 µm.

**(E)** Western blot using anti-myc antibodies on total protein extracts from cells expressing *ipl1-5as-*myc treated 60 h with DMSO or 50 µM NA-PP1 after quiescence entry. Ade13 was used as a loading reference.

**(F)** Nuclear MT length in cells of the indicated genotypes grown 48 h at 37 °C or 96 h at 25 °C, treated 15 min or not with 30 µg/ml Noc. Same legend as in (B). Images of representative cells are shown, bar is 2 µm.

**(G)** Distribution of Bim1-3GFP intensity as a function of mTQZ-Tub1 intensity in individual Q-nMT bundles (green: phase I – 7 h after glucose exhaustion, orange: phase II – 26 h after glucose exhaustion and red: phase III – 50h after glucose exhaustion). Each circle represents an individual Q-nMT bundle.

**(H)** Viability (methylene blue staining) after the indicated time in quiescence (left graph), capacity to form colony after single live cell micro-manipulation after 12 days in quiescence (right graph). Pictures of representative plates 3 d after cell micro-manipulation are shown.

**(I)** Western blot using anti-GFP antibodies on total protein extracts from proliferating or 4-days-old cells expressing the indicated GFP fusion protein. Ade13 and Act1 were used as loading controls.

**(J)** Percentage of cells of the indicated genotype displaying a MT star-like array. Each circle represents the % obtained for an independent culture (n > 200). Mean and SD are indicated.

**(K)** WT cells (6 d) expressing mRuby-Tub1 (red) and Nuf2-GFP (green) were transferred onto an agarose pad. Q-nMT bundles were cut using a pulsed laser (pink dashed line). White arrowheads point to dynamic cMTs. Time is in min, bar is 2 µm.

**(L)** Proliferating cells expressing mRuby-Tub1 (red) and Dad2-GFP (green) were transferred onto an agarose pad. Anaphase spindles were cut and the behavior of the released fragments was followed. Time is in min, bar is 2 μm.

**(M)** Nuclear MT length in cells of the indicated genotypes, grown for 3 d at 25 °C, shifted to 37 °C or 25° C for 48 h, and treated 15 min or not with 30 µg/ml Noc. Same legend as in (B). Images of representative cells are shown, p-value from unpaired t-test is indicated, bar is 2 µm.

**Supplementary Figure 4**

**(A)** Q-nMT bundle length as a function of Q-nMT bundle width for individual WT cells (grey) and *kar3Δ* cells (red) 90 h after glucose exhaustion.

**(B)** WT cells expressing mTQZ-Tub1 (red) and Cin8-GFP (green) were imaged at the indicated times after glucose exhaustion. Cells expressing only mTQZ-Tub1 (left panel) testify for the absence of fluorescence leak from the mTQZ signal into the GFP channel. Bar is 2μm.

**(C)** Fluorescence intensity along Q-nMT bundles in WT (black) and *cin8Δ* cells expressing mTQZ-Tub1, treated (green) or not (red) with Noc. Each line is an individual cell, the bold lines are the mean (n > 20).

**(D)** Morphometric Q-nMT bundle properties distribution in WT (grey) and *cin8Δ* (green) cells expressing mTQZ-Tub1. Each circle is an individual cell.

**(E)** Fluorescence intensity along Q-nMT bundles in WT (grey, n > 50), *kip1Δ* (red, n > 50), *bim1Δ* (green, n > 50), and *kip1Δ bim1Δ* (blue, n > 30) cells (4 d). Each line is an individual cell, the bold lines are the mean.

**(F)** Nuclear MT length distribution in *kip2Δ* cells expressing mTQZ-Tub1 treated 15 min (blue dots) or not (black dots) with 30 µg/ml Noc. For a comparison, the mean WT length is indicated as dashed line. Unpaired t-test p-values are indicated.

**Supplementary Figure 5**

**(A)** Representative 5-day-old cells (left and right) expressing mTQZ-Tub1 (red) and Nuf2-GFP (green) exiting quiescence on a YPDA agarose pad. Blue arrowhead: SBP; white arrows Nuf2-GFP clusters. Time is in min, bar is 1µm.

**(B)** WT and *kip3Δ* cells (5 d) expressing Nuf2-GFP were re-fed on an YPDA microscope pad.

Each line corresponds to an individual cell. Data were presented either (left) with the time set to zero at the onset MT depolymerisation or (right) at the onset of SPB separation (red dash line).

**(C)** Western blot using GFP antibodies on total protein extracts from WT cells expressing Kip3-GFP grown for the indicated time. Sac6 was used as a loading control.

**(D-E)** Nuclear Q-nMT bundle length in cells of the indicated genotype expressing mTQZ-Tub1, grown for 4 d and imaged at the indicated time after refeeding. For the rapamycin experiment in (D), WT cells were pre-incubated 1 h with rapamycin (1 μg/mL), and then re-fed in presence of the drug. For thermo-sensitive strains in (E), cells were grown 4 d at 25 °C, shifted 1 h at 37 °C, and then transferred to new medium at 37 °C to trigger quiescence exit. Measurements were done after a 15 min Noc treatment (30 µg/ml) to remove dynamic cMTs. Each circle represents an individual cell. Mean and SD are indicated.

**(F-G)** Nuclear Q-nMT bundle length in cells of the indicated genotype expressing mTQZ-Tub1 grown for the indicated time at 25 °C, shifted to 37 °C and re-fed. In G, measurements were done after a 15 min Noc treatment (30 µg/ml) to remove dynamic cMTs. Each circle represents an individual cell. Mean and SD are indicated.

**(H)** 3 individual experiments in which nuclear MT length was measured in 4-day-old *kip3Δshe1Δ* cells expressing mTQZ-Tub1 upon quiescence exit (n > 100). P-values from unpaired t-test are shown. Each circle represents an individual cell. Mean and SD are indicated.

**(I)** Percentage of cells with separated SPBs upon quiescence exit on YPDA agarose pad (N = 4, n > 200, left panel). Right: proliferation of WT and *kip3Δshe1Δ* strains in YPDA (OD 600nm as function of time).

**(J)** Q-nMT bundle length distribution in WT (grey) or *kip3Δshe1Δ* (green) cells with unseparated SPBs, 240 min after quiescence exit.

**(K)** Actin (red, phalloidin-staining) in WT and *kip3Δshe1Δ* cells expressing mTQZ-TUB1 (green) at the indicated time upon quiescence exit. Bar is 2μm.

## Notes

### Competing Interest Statement

The authors have declared no competing interest.

### Summary of Updates

Some phrases/words have been changed for better understanding.

