## Supplementary figures and images for "A stable microtubule bundle formed through an orchestrated multistep process controls quiescence exit"

### Sup Figure 1-5

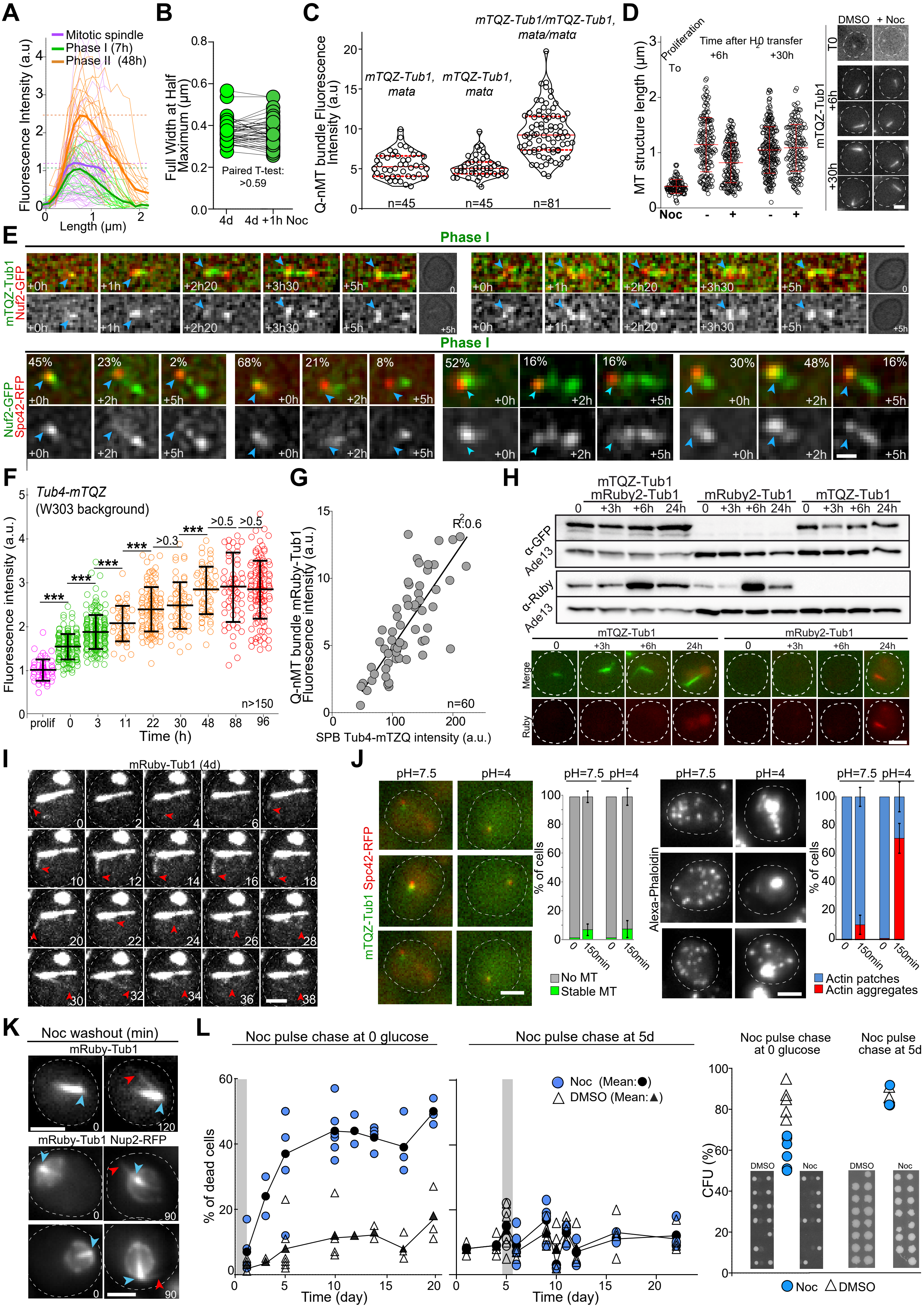

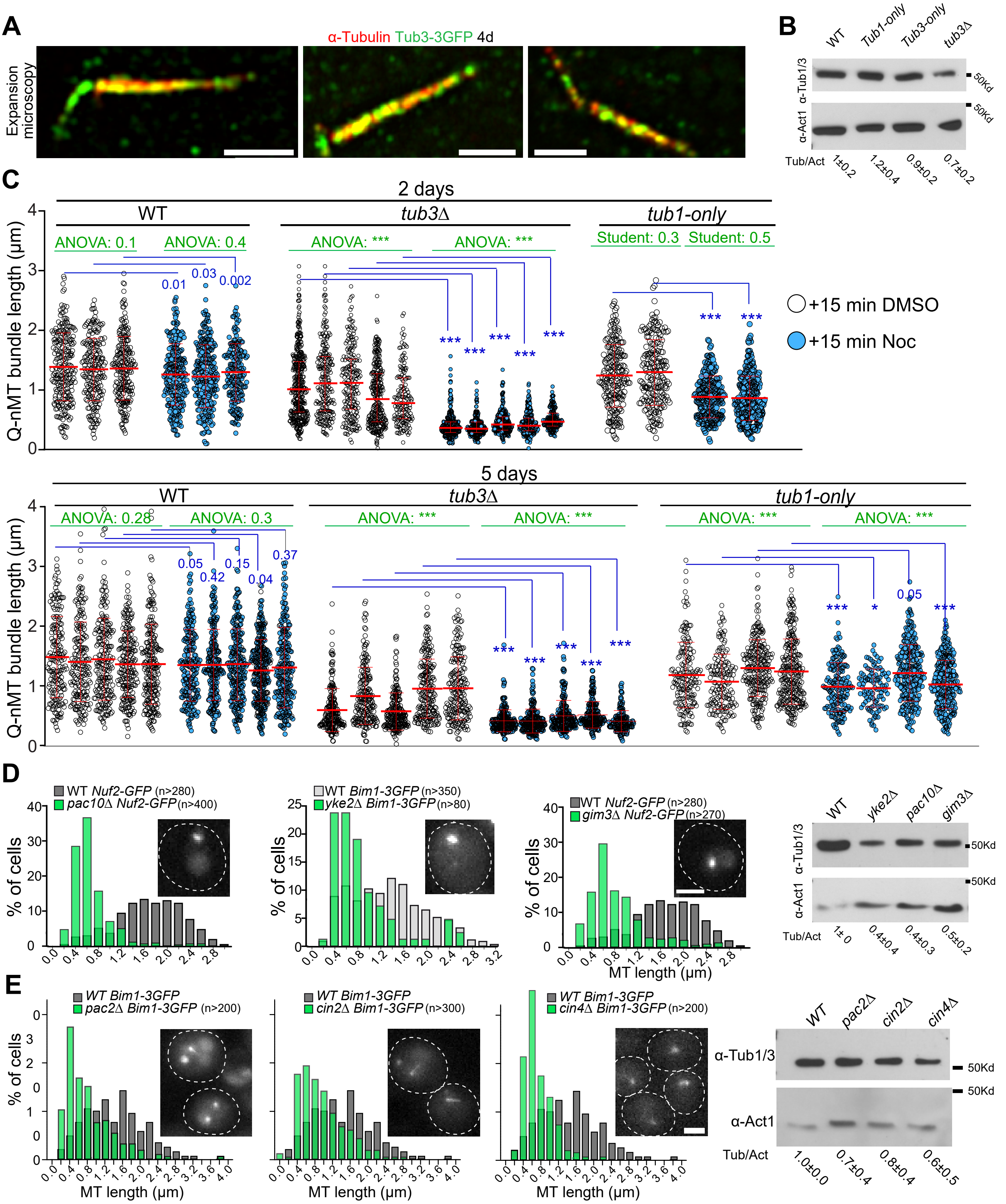

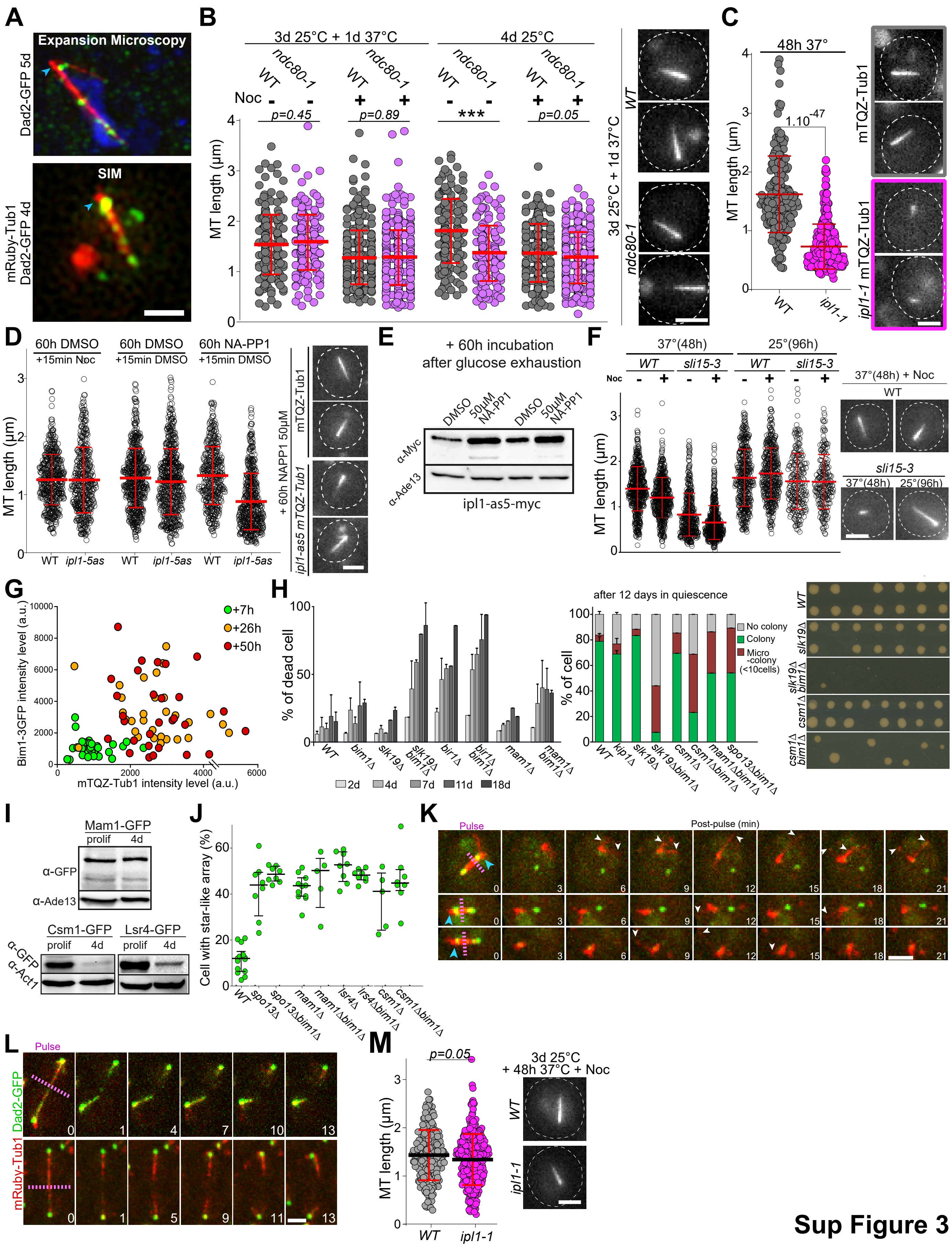

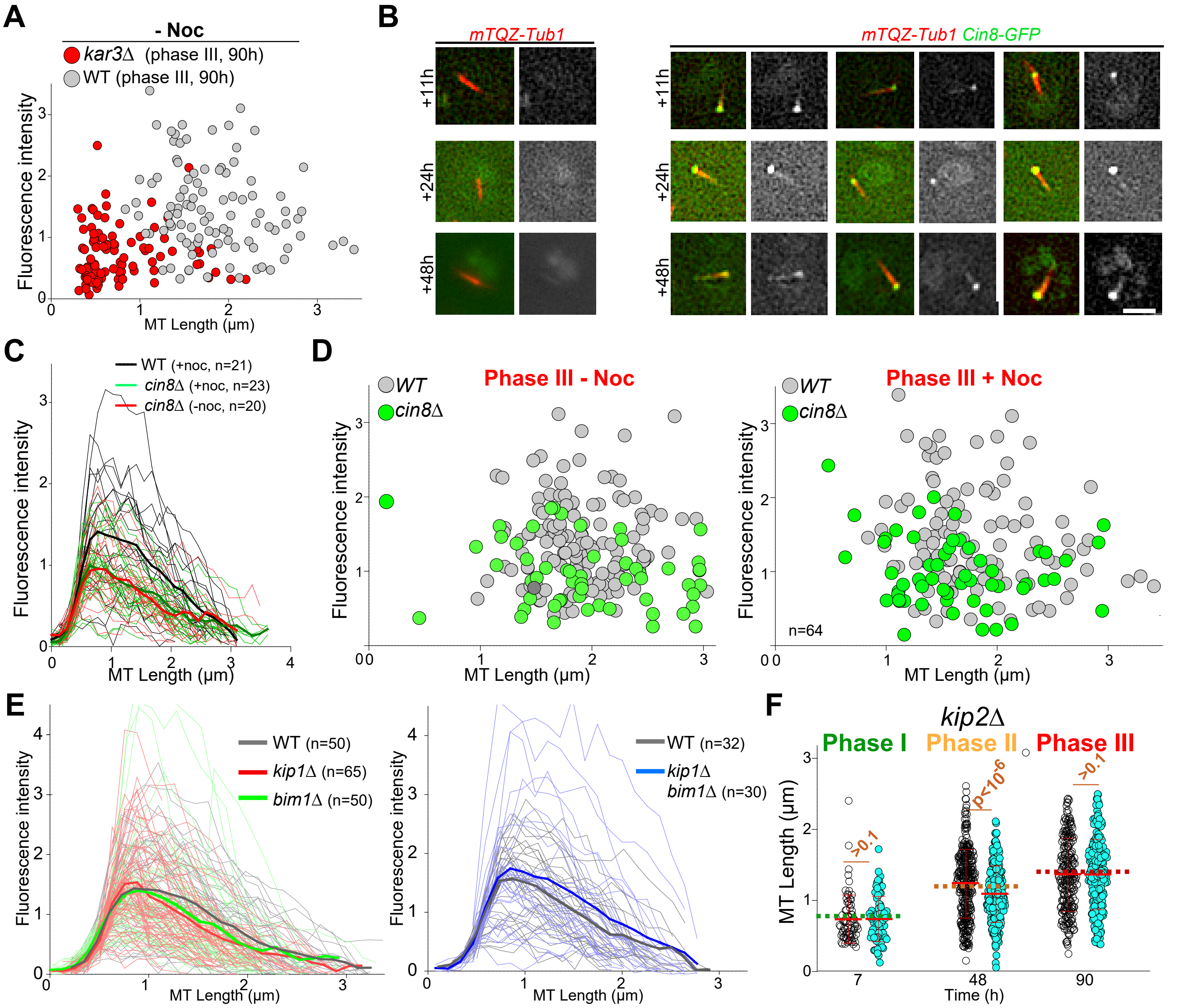

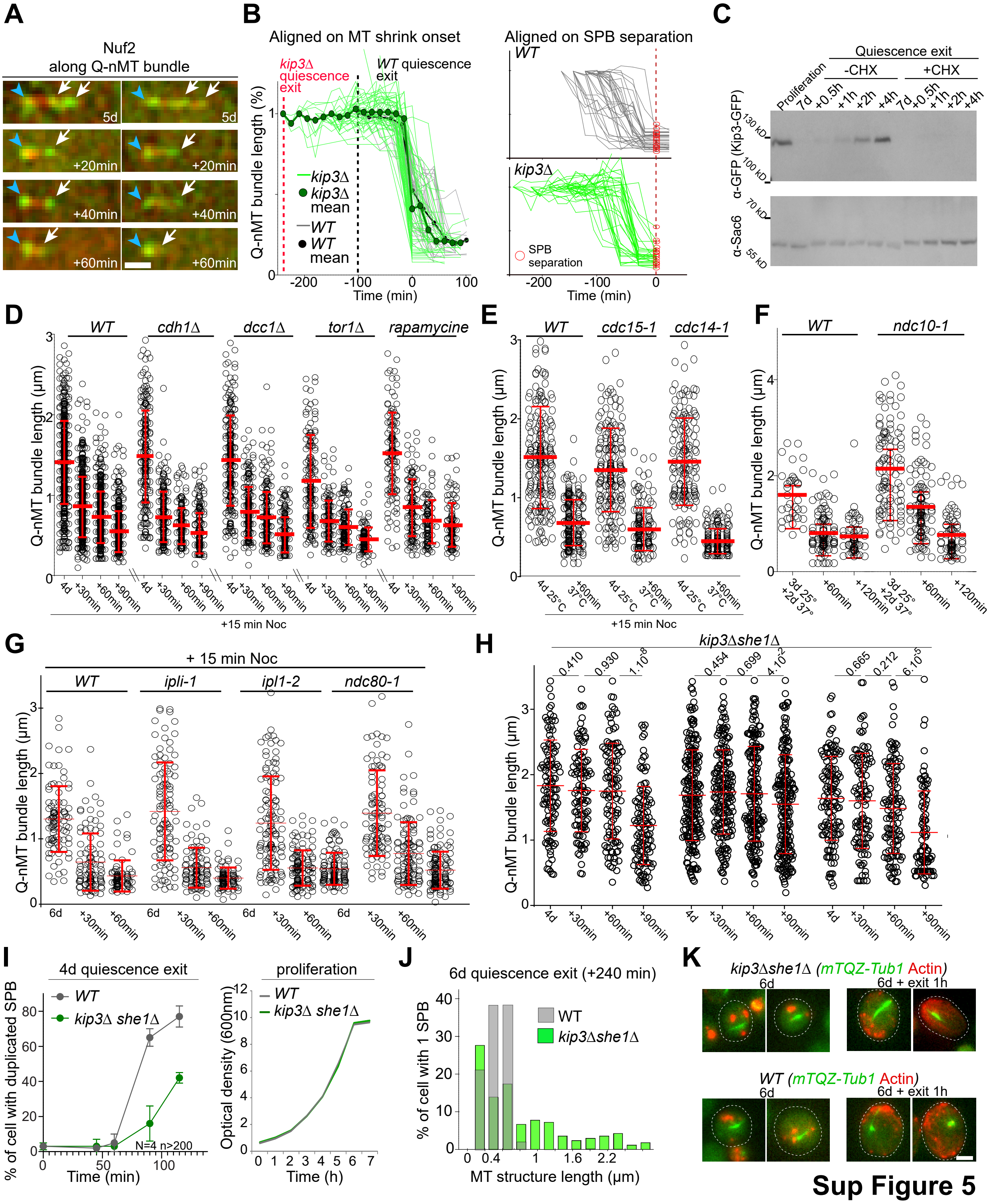
